# The sulfonylpiperazine MMV020291 prevents red blood cell invasion by the malaria parasite *Plasmodium falciparum* through interference with actin-1/profilin dynamics

**DOI:** 10.1101/2022.09.29.510018

**Authors:** Madeline G. Dans, Henni Piirainen, William Nguyen, Sachin Khurana, Somya Mehra, Zahra Razook, Sujaan Das, Molly Parkyn Schneider, Thorey K. Jonsdottir, Mikha Gabriela, Maria R. Gancheva, Christopher J. Tonkin, Vanessa Mollard, Christopher Dean Goodman, Geoffrey I. McFadden, Danny W. Wilson, Alyssa E. Barry, Brendan S. Crabb, Tania F. de Koning-Ward, Brad E. Sleebs, Inari Kursula, Paul R. Gilson

## Abstract

With emerging resistance to frontline treatments, it is vital that new antimalarial drugs are identified to target *Plasmodium falciparum*. We have recently described a compound, MMV020291, as a specific inhibitor of red blood cell invasion, and have generated analogues with improved potency. Here, we identify actin and profilin as putative targets of the MMV020291 series through resistance selection and whole genome sequencing of three MMV020291 resistant populations. This revealed three non-synonymous single nucleotide polymorphisms in two genes; two in *profilin* (N154Y, K124N) and a third one in *actin-1* (M356L). Using CRISPR-Cas9, we engineered these mutations into wildtype parasites which rendered them resistant to MMV020291. We demonstrate that MMV020291 reduces actin polymerisation that is required by the merozoite stage parasites to invade red blood cells. Additionally, the series inhibits the actin-1 dependent process of apicoplast segregation, leading to a delayed death phenotype. *In vitro* co-sedimentation experiments using recombinant *P. falciparum* actin-1 and profilin proteins indicate that potent MMV020291 analogues amplify the actin-monomer sequestering effect of profilin, thereby reducing the formation of filamentous actin. Altogether, this study identifies the first compound series targeting the actin-1/profilin interaction in *P. falciparum* and paves the way for future antimalarial development against the highly dynamic process of actin polymerisation.

## Introduction

Malaria is a devastating parasitic disease that caused ∼ 627 000 deaths in 2020, an upward trend from the previous year’s figure of 558 000 due to COVID-19-related service disruption (World Health Organization, 2020). The majority of these deaths were a result of infection with *Plasmodium falciparum*, which causes widespread disease across Sub-Saharan Africa. The alarming upward trend in deaths combined with the spread of mutations conferring resistance to frontline antimalarials (Dondorp et al., 2009; Fairhurst and Dondorp, 2016; Uwimana et al., 2020), highlights the urgent need to find new compounds with unique mechanisms of action (MoA) to combat this deadly parasite.

The red blood cell (RBC) stage of *Plasmodium* infection within the human host leads to the exponential growth of the parasite and the symptoms of the disease. Within RBCs, parasites develop within a parasitophorous vacuole (PV) in a series of stages from rings to trophozoites and finally schizonts. During schizogony daughter merozoites are formed, which eventually egress from the RBC to re-infect new RBCs. The invasion of RBCs by merozoites is a complex and finely-tuned process that involves a multitude of unique signalling cascades and protein-protein interactions (Cowman et al., 2017). These events must occur within minutes of a merozoite egress to allow successful RBC internalisation and, therefore, this process represents a prime opportunity against which to develop therapeutics (Boyle et al., 2010; Burns et al., 2019; Gilson and Crabb, 2009; Weiss et al., 2015). Administering an invasion inhibitor in combination with a drug targeting intracellular parasite processes would span the entire asexual blood stage of infection and could effectively prevent parasite proliferation (Burns et al., 2019).

One unique process required for parasite invasion of RBCs is the engagement of an actomyosin motor complex, termed the glideosome, a mechanism that is shared between apicomplexan parasites. In the glideosome, a single-headed class XIV myosin A, MyoA, is tethered to the inner membrane complex of the merozoite via its two light chains and several glideosome-associated proteins (Frénal et al., 2010; Green et al., 2006; Green et al., 2017). MyoA produces the force required for gliding motility by walking along filamentous actin (Blake et al., 2020; Das et al., 2021; Robert-Paganin et al., 2019; Vahokoski et al., 2022). Actin filaments, in turn, are linked to surface exposed adhesin proteins via the glideosome associated connector (Jacot et al., 2016). A ring of adhesions, termed the tight junction, is then formed between the apical tip within the merozoite and the RBC and the force produced by the MyoA power stroke, propels the merozoite into the RBC through the established tight junction (Cowman and Crabb, 2006; Keeley and Soldati, 2004). This movement translocates the tight junction from the apical to the posterior end of the parasite and results in the RBC membrane enveloping the merozoite that later forms the PV (Cowman et al., 2017; Dobrowolski et al., 1997; Gonzalez et al., 2009).

The generation of filamentous actin (F-actin) is required for both gliding motility and RBC invasion by the parasite (Douglas et al., 2018; Yahata et al., 2021). The transition between globular actin (G-actin) and F-actin, referred to as actin treadmilling, is a highly regulated process that involves the hydrolysis of G-actin-ATP to G-actin-ADP and inorganic phosphate (P_i_) at the barbed end to stabilise the growing filament (Fujiwara et al., 2002; Korn et al., 1987; Wegner, 1976). For filament disassembly and G-actin turnover, the release of P_i_ destabilises the F-actin, resulting in the disassociation of G-actin-ADP (Carlier, 1990; Carlier and Pantaloni, 1986). This actin treadmilling process is tightly regulated by a plethora of actin-binding regulatory proteins, many of which are absent in apicomplexan parasites (Baum et al., 2006; Das et al., 2021). However, one key protein present across eukaryotes from Apicomplexa to Opisthokonts is profilin, a sequester of G-actin (Carlsson et al., 1977). The role of profilin is to maintain a pool of polymerizable actin monomers, and to catalyse the exchange of ADP to ATP in the monomers (Mockrin and Korn, 1980; Witke, 2004). Profilin then delivers these G-actin-ATP to formin, a nucleator and processive capper that binds to the barbed end of the actin filament (Courtemanche, 2018; Funk et al., 2019; Goode and Eck, 2007).

Profilin is essential for merozoite invasion of RBCs (Kursula et al., 2008; Pino et al., 2012) and is required for efficient sporozoite motility (Moreau et al., 2017; Moreau et al., 2021). Apicomplexan profilin has diverged markedly from higher eukaryotes (Kursula et al., 2008), with low sequence identity (16%) retained between *P. falciparum* profilin (PfPFN) (PF3D7_0932200) and *Homo sapiens* profilin I (HsPFNI) sequences. This is illustrated by a unique arm-like β-hairpin insertion in apicomplexan profilin that spans from residues 57-74 (Kursula et al., 2008). This unique sequence is essential for actin binding; mutations in this region hinder monomer sequestration *in vitro* and impact sporozoite motility (Moreau *et al*., 2017). Within apicomplexan, PfPFN has also diverged from *Toxoplasma gondii* profilin (TgPFN), with the former acquiring a further acidic loop extension located in residues 40-50 (Kursula et al., 2008).

In comparison, actin is more conserved between apicomplexans and higher eukaryotes, however the apicomplexan actins are among the most diverged actins in eukaryotes. *P. falciparum* encodes two actin isoforms; actin-1 (PfACT1) (PF3D7_1246200) and actin-2 (PfACT2) (PF3D7_1412500), the latter of which is only expressed in the sexual stages (Wesseling et al., 1989). PfACT1 has a high degree of sequence identity (93%) with the single actin gene in *T. gondii* (Dobrowolski et al., 1997) and shares an 82% identical sequence with the human cytosolic β actin (Baum et al., 2006; Das et al., 2021; Wesseling et al., 1989).

In apicomplexan parasites, actin polymerisation has been difficult to study because of the short length and instability of the actin filaments (Das et al., 2021; Kumpula et al., 2019; Schmitz et al., 2005). Despite this, actin polymerisation has been shown to be essential in many phases of the lifecycle, including intracellular replication, host cell egress (only in *T. gondii*), motility and host cell invasion (Das et al., 2021; Jacot et al., 2016; Periz et al., 2017; Tardieux and Baum, 2016). Of these, host cell invasion is the best-characterised actin-dependent process (reviewed in (Tardieux and Baum, 2016). In *P. falciparum*, naturally occurring compounds such as cytochalasins D and B, latrunculins, phalloidin and jasplakinolide have been used to study the complex actin regulation in merozoite invasion (Baum et al., 2006; Johnson et al., 2016; Miller et al., 1979; Pospich et al., 2017; Varghese et al., 2020; Weiss et al., 2015). These compounds interfere with actin treadmilling by affecting the polymerisation and depolymerisation of actin through various MoA. For example cytochalasins bind to the barbed end to prevent polymerisation and can also prevent G-actin disassociation from F-actin (Shoji et al., 2012), latrunculins prevent sequestration of the G-actin subunits (Morton et al., 2000) and phalloidin and jasplakinolide stabilise F-actin by preventing the release of P_i_ from G-actin-ADP subunits (Pospich et al., 2020). Apart from the recently developed latrunculins that have greater selectivity (Varghese et al., 2020), these naturally occurring compounds remain biological tools rather than antimalarial candidates due to their cytotoxicity.

Recently, we identified a compound MMV020291 (MMV291) from the Medicines for Malaria Pathogen Box as an inhibitor of *P. falciparum* merozoite invasion of RBCs (Dans et al., 2020). We further explored the drug development potential of the compound by defining the structure activity relationship (SAR) and generated analogues with improved potency, whilst maintaining compound selectivity and invasion blocking activity (Nguyen et al., 2021). Here, through *in vitro* resistance selection, whole genome analysis and reverse genetics we show that the putative targets of MMV291 are PfPFN and PfACT1. We further explore the MoA of the MMV291 which is the first reported compound series targeting the actin-profilin complex in *P. falciparum*.

## Results

### MMV291 resistant parasites contain mutations in *actin-1* and *profilin*

To select for parasite resistance against our lead molecule MMV291 (Fig. 1A), five populations of 10^8^ *P. falciparum* 3D7 parasites were exposed to 10 µM (∼10×EC_50_) of the compound until new ring stage parasites were no longer observed by Giemsa-stained blood smears. The drug was removed, and parasites were allowed to recover. This drug on and off selection was performed for three cycles before parasite resistance was evaluated in a 72-hour lactate dehydrogenase (LDH) growth assay (Makler and Hinrichs, 1993). This revealed three MMV291-selected populations that demonstrated an 8 to 14-fold increase in EC_50_ (Fig. S1). These resistant populations (B, C and D) were cloned out by limiting dilution and two clones from each parent line were tested in an LDH assay, indicating resistance was heritable (Fig. 1B). Genomic DNA was extracted from these resistant clones, along with a parental 3D7 reference strain, and whole genome sequencing was performed using the Oxford Nanopore MinION platform (Razook et al., 2020). Here, a minimum of 10x depth coverage with 70% of the reads to support the called allele was required for verification (Razook et al., 2020).

**Figure 1.**
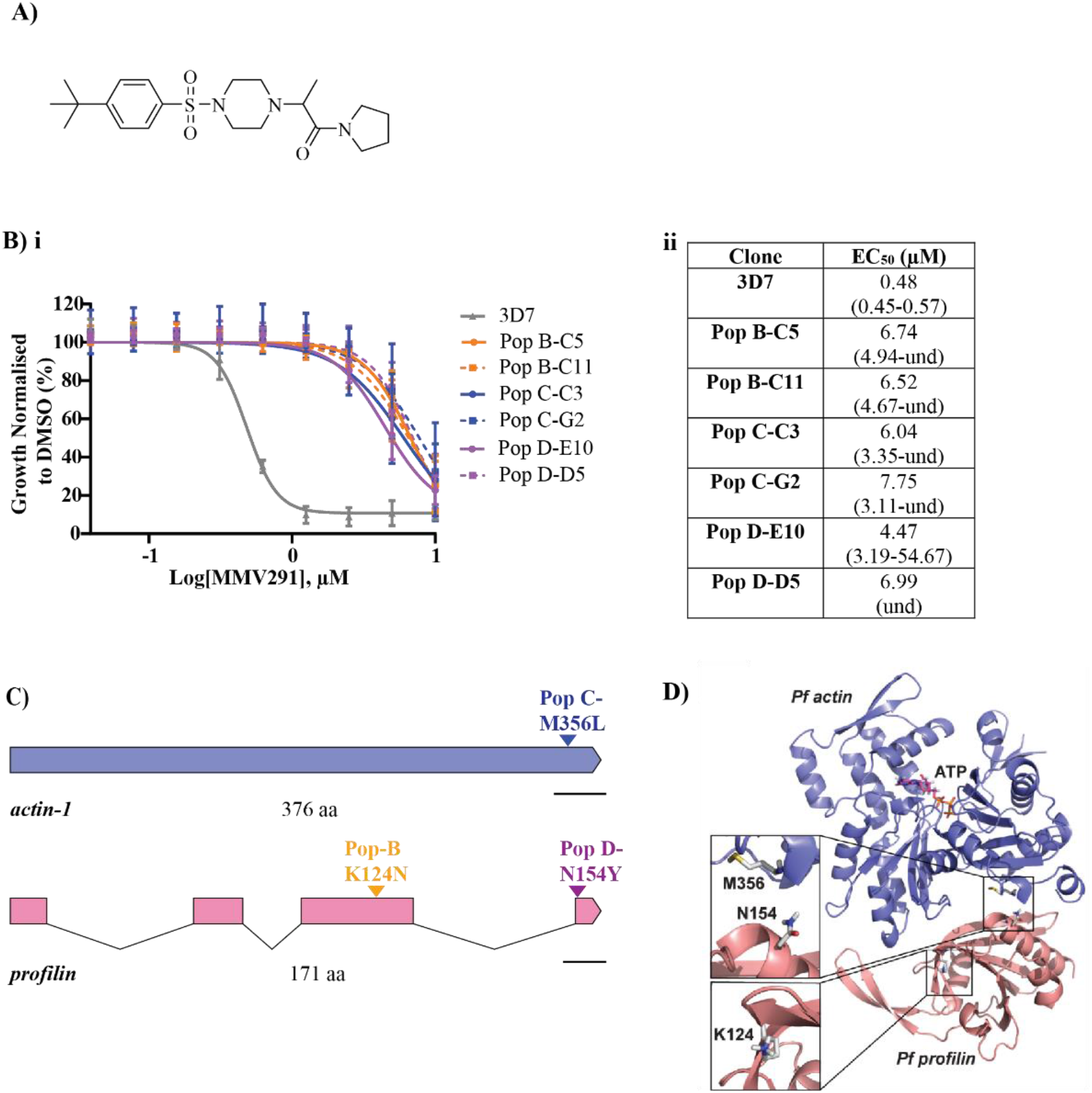
Resistance selection and whole genome sequencing reveal *actin-1* and *profilin* as putative targets of MMV291A) Chemical structure of MMV291. **B)** i Drug cycling on and off for three cycles and subsequent cloning out of parental lines resulted in two clones from three populations (Pop B, C and D) that maintained stable resistance to MMV291 in a 72-hour growth assay. Growth has been normalised to that of parasites grown in 0.1% DMSO with error bars representing the standard deviation of three biological replicates. **ii** EC_50_ values derived from nonlinear regression curves in GraphPad Prism with 95% confidence intervals shown in brackets. und =undefined. **C)** Genome sequencing of the MMV291-resistant parasites revealed a different non-synonymous single nucleotide polymorphism (SNP) shared by the clonal lines across two related proteins; Population D and B contained K124N and N154Y mutations in profilin (PF3D7_0932200), respectively, whilst Population C contained a M356L mutation in actin-1 (PF3D7_1246200). Scale bar indicates 100 base pairs. **D)** The positions of the resistance mutations were mapped onto the X-ray structures of *P. falciparum* actin-1 (purple) (PDB: 6I4E) (Kumpula et al., 2019) and *P. falciparum* profilin (pink) (PDB: 2JKG) (Kursula et al., 2008) which revealed PFN(N154Y) and ACT1(M356L) lie on either side of the proteins’ binding interfaces. PFN(K124N) resides on the opposing side of profilin. In this case, the X-ray structures of *Oryctolagus cuniculus* actin and human profilin (PDB: 2PBD) (Ferron et al., 2007) were utilised as a template to spatially align the two parasite proteins.

Across the six clones of MMV291-resistant parasites from three populations, there were a total of 18 non-synonymous single nucleotide polymorphisms (SNPs) identified in 16 genes with no other gene variants found (Table 1). Of these SNPs, three were present in related genes across all resistant isolates. One of these SNPs was located in chromosome (Chr) 12:1921849 within the gene encoding *actin-1* (PF3D7_1246200), resulting in a M356L mutation in population C clones. The other two SNPs both occurred in the gene encoding *profilin* (PF3D7_0932200), located in Chr 9:1287853 and 1288316, resulting in a K124N and N154Y mutation in population B and D clones, respectively (Fig. 1C, Table 1).

**Table 1.** Resistant genomes pooled variant summary. Table depicts genes in which non-synonymous single nucleotide polymorphisms (SNPs) were identified that passed quality filtration and were detected in at least one resistant clone. Highlighted rows indicate profilin and actin-1 as the putative targets of MMV291.

| Chromosome | Position | Ref | Alt | Effect | Gene_ID | Gene_Description | Codon<br>Change | AA_<br>Change | WT_3<br>D7_C9 | WT_3<br>D7 | Pop<br>B_C11 | Pop<br>B_C5 | Pop<br>C_C3 | Pop<br>C_G2 | Pop<br>D_D5 | Pop<br>D_E10 |
| --- | --- | --- | --- | --- | --- | --- | --- | --- | --- | --- | --- | --- | --- | --- | --- | --- |
| Pf3D7_09_v3 | 1287853 | A | T | nonsynonymous | PF3D7_0932200 | *profilin* | AAA/AAT | K124N | . | . | ALT1 | ALT1 | . | . | . | . |
| Pf3D7_09_v3 | 1288316 | A | T | nonsynonymous | PF3D7_0932200 | *profilin* | AAT/TAT | N154Y | . | . | . | . | . | . | ALT1 | ALT1 |
| Pf3D7_12_v3 | 1921849 | A | T | nonsynonymous | PF3D7_1246200 | *actin 1* | ATG/TTG | M356L | . | . | . | . | ALT1 | ALT1 | . | . |
| Pf3D7_06_v3 | 554517 | T | G | nonsynonymous | PF3D7_0613600 | *conserved Plasmodium*<br>*protein, unknown function* | TTT/GTT | F21V | . | . | ALT1 | ALT1 | . | . | . | . |
| Pf3D7_03_v3 | 547075 | G | T | nonsynonymous | PF3D7_0313400 | *conserved Plasmodium*<br>*protein, unknown function* | CAA/AAA | Q370K | . | . | . | . | . | . | . | ALT1 |
| Pf3D7_04_v3 | 682224 | G | A | nonsynonymous | PF3D7_0415300 | *cdc2-related protein kinase 3* | GAA/AAA | E139K | . | . | . | . | . | . | . | ALT1 |
| Pf3D7_06_v3 | 524255 | C | T | nonsynonymous | PF3D7_0612600 | *cytoplasmic tRNA 2-thiolation*<br>*protein 1, putative* | GGA/GAA | G284E | . | . | . | . | ALT1 | . | . | . |
| Pf3D7_07_v3 | 909422 | C | A | nonsynonymous | PF3D7_0721000 | *conserved Plasmodium*<br>*membrane protein, unknown*<br>*function* | CAA/AAA | Q1711K | . | . | . | . | . | . | . | ALT1 |
| Pf3D7_11_v3 | 1266495 | C | T | nonsynonymous | PF3D7_1132600 | *pre-mRNA-splicing factor 38A,*<br>*putative* | GAA/AAA | E306K | . | . | . | . | . | . | . | ALT1 |
| Pf3D7_12_v3 | 513882 | C | A | nonsynonymous | PF3D7_1211600 | *lysine-specific histone*<br>*demethylase 1, putative* | ACA/AAA | T1187K | . | . | . | . | . | . | . | ALT1 |
| Pf3D7_12_v3 | 1022182 | G | T | nonsynonymous | PF3D7_1225100 | *isoleucine--tRNA ligase,*<br>*putative* | GTT/TTT | V1114F | . | . | . | . | . | . | . | ALT1 |
| Pf3D7_12_v3 | 1381681 | C | T | nonsynonymous | PF3D7_1233400 | *conserved Plasmodium*<br>*membrane protein, unknown*<br>*function* | AGA/AAA | R233K | . | . | . | . | . | . | . | ALT1 |
| Pf3D7_13_v3 | 1004824 | G | T | nonsynonymous | PF3D7_1324300 | *conserved Plasmodium*<br>*membrane protein, unknown*<br>*function* | CCA/CAA | P4827Q | . | . | . | . | ALT1 | . | . | . |
| Pf3D7_13_v3 | 1810063 | G | A | nonsynonymous | PF3D7_1345100 | *thioredoxin 2* | GCT/GTT | A84V | . | . | . | . | ALT1 | . | . | . |
| Pf3D7_14_v3 | 271754 | C | T | nonsynonymous | PF3D7_1407600 | *conserved Plasmodium*<br>*protein, unknown function* | GAA/AAA | E1788K | . | . | . | . | . | . | . | ALT1 |
| Pf3D7_14_v3 | 404507 | G | T | nonsynonymous | PF3D7_1410300 | *WD repeat-containing protein,*<br>*putative* | CAA/AAA | Q4186K | . | . | . | . | . | . | . | ALT1 |
| Pf3D7_14_v3 | 413594 | C | T | nonsynonymous | PF3D7_1410300 | *WD repeat-containing protein,*<br>*putative* | GAA/AAA | E1157K | . | . | . | . | . | . | . | ALT1 |
| Pf3D7_14_v3 | 1464591 | C | A | nonsynonymous | PF3D7_1436100 | *conserved Plasmodium*<br>*membrane protein, unknown*<br>*function* | AAC/AAA | N1068K | . | . | . | . | . | . | . | ALT1 |

A homology model of the binding of *P. falciparum* actin-1 (PfACT1) and profilin (PfPFN) was created using the binding of *Orytolagus cuniculus* actin to *H. sapiens* profilin (Ferron et al., 2007; Kumpula et al., 2019; Kursula et al., 2008) (Fig. 1D). This indicated that PfACT1(M356) and PfPFN(N154) were located at the binding interface between the two proteins, whilst PfPFN(K124) was orientated away, on the opposite side of PfPFN. Despite the close proximity of these two SNPs to the binding site between the two proteins, the resistant parasites did not exhibit an associated fitness cost *in vitro* (Fig. S2), indicating these amino acid changes are well-tolerated and may not be essential for actin-1 binding to profilin.

### Mutations in *actin-1* and *profilin* mediate resistance to MMV291 in wildtype parasites

To confirm that the chemically induced PfPFN(N154Y), PfPFN(K124N) and PfACT1(M356L) mutations were responsible for resistance to MMV291, we employed reverse genetics to introduce each mutation into wildtype parasites. This was achieved using a CRISPR-Cas9 gene editing system whereby homology regions were designed to both the 5’ and 3’ flanks of the desired loci. The 5’ homology flank encompassed a synthetic recodonised region containing the wildtype or non-synonymous drug resistant mutations and synonymous shield mutations to prevent re-cleavage with Cas9 after recombination into the desired loci (Fig. 2Ai). To direct Cas9 cleavage, a homologous synthetic guide RNA (gRNA) was mixed with a tracrRNA and recombinant Cas9 enzyme and electroporated into blood stage parasites with the donor plasmid (McHugh et al., 2021). After chromosomal integration was selected with WR99210, viable parasites for both the mutant and wildtype parasites were confirmed to contain the donor cassette using integration PCRs (Fig. 2Aii). These PCR products were sequenced and confirmed to contain the corresponding MMV291-resistant alleles (Fig. S3). Next, the modified lines were tested in a LDH growth assay against MMV291 which showed an 11 to 18-fold increase in EC_50_ in the introduced mutant lines compared to their wildtype counterparts (Fig. 2B). This indicated that PfPFN(K124N), PfPFN(N154Y) and PfACT1(M356L) were responsible for the chemically induced resistance by MMV291, suggesting these proteins are the putative targets of the compound.

**Figure 2.**
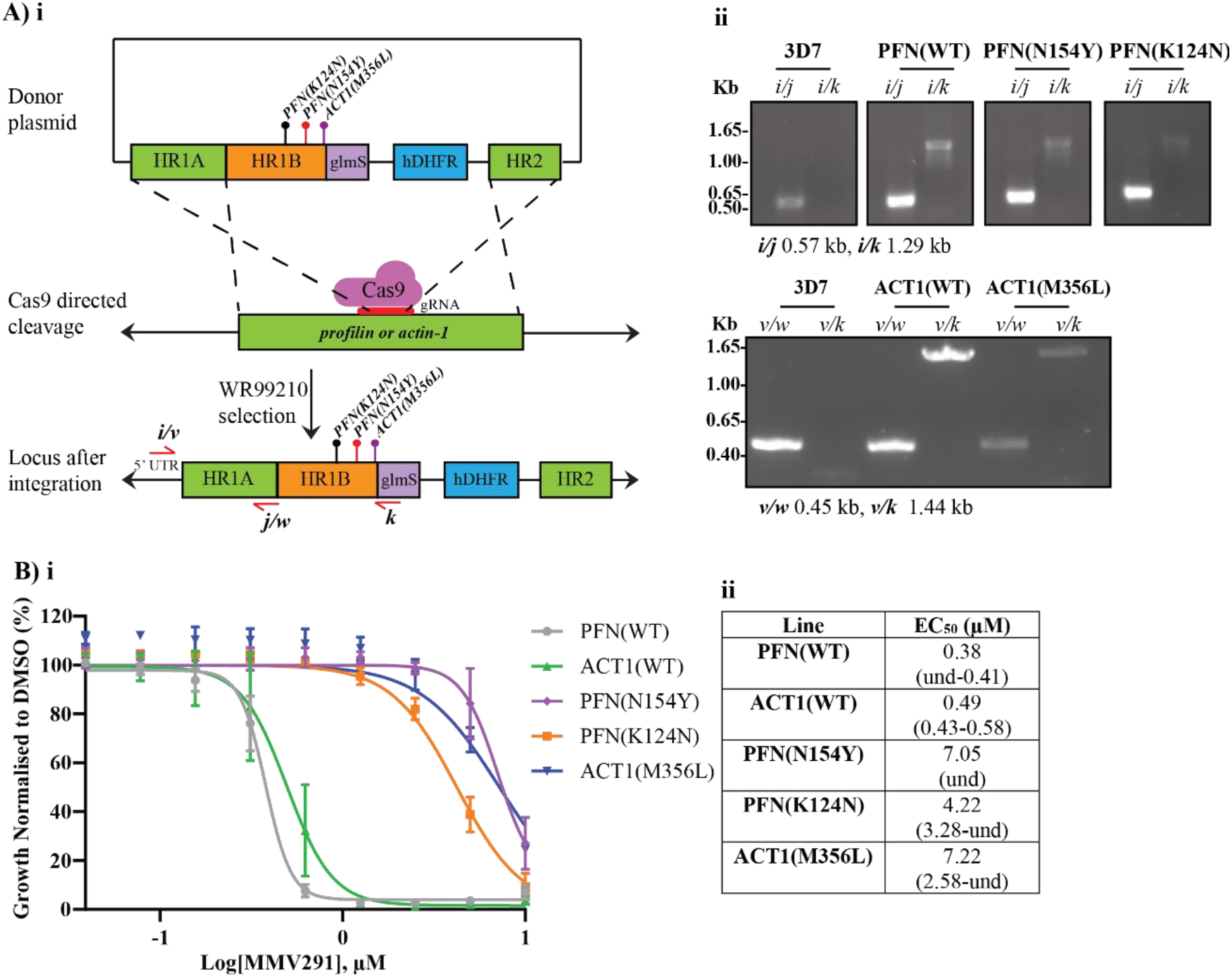
Introduction of the SNPs in *profilin* and *actin-1* into 3D7 parasites mediates resistance to MMV291A) i. Strategy to create the donor plasmid to introduce PFN(N154Y), PFN(K124N) and ACT1(M356L) SNPs into 3D7 parasites. Homology regions (HR) were designed to the 5’ flank (HR1) and 3’flank (HR2) whereby HR1 was made up of the endogenous genes’ sequence (HR1A) and recodonised fragments (HR1B), encompassing the resistant mutation alleles. A synthetic guide RNA (gRNA) was designed for either *profilin* or *actin-1* to direct Cas9 to the cleavage site and induce double crossover homologous recombination. WR99210 was used to select for integrated parasites via the human hydrofolate reductase (hDHFR). **ii** Integration into the *profilin* or *actin-1* locus was validated whereby a 5’ UTR primer (a) was used in combination with a primer located in the glmS region (c). **B) i** Integrated parasites were tested in a 72-hour LDH growth assay which revealed the resistant mutations conferred resistance against MMV291 and confirmed the profilin and actin-1 proteins as putative targets of the compound. Growth has been normalised to that of parasites grown in 0.1% DMSO and error bars indicate the standard deviation of three biological replicates. **ii** EC_50_ values derived from nonlinear regression curves in GraphPad Prism with 95% confidence intervals shown in brackets. und =undefined.

To date, in *P. falciparum*, the dynamics of actin polymerisation has been explored with naturally occurring compounds that bind to various regions within the actin-1 protein (Baum et al., 2008; Johnson et al., 2016; Miller et al., 1979; Pospich et al., 2017; Pospich et al., 2020; Varghese et al., 2020; Weiss et al., 2015) (Fig. S4). Targeting the actin-binder profilin, however, presents a novel mechanism to interfere with this essential parasite process. This novel MoA of MMV291 was confirmed by the lack of cross-resistance between the chemically induced MMV291-resistant parasites and cytochalasin D (CytD) and jasplakinolide in a 72-hour LDH growth assay (Fig. S5). Furthermore, despite the highly conserved sequence of actin-1 in *H. sapiens* and *P. falciparum* (Fig. S6) and that actin dynamics in RBCs have previously been shown to be perturbed by actin inhibitors (Gokhin and Fowler, 2016; Gokhin et al., 2015), RBCs that had been pre-treated with MMV291 displayed normal levels of merozoite invasion, indicating this compound ins not targeting host actin (Fig. S7). Altogether, this data indicates that MMV291 has an alternative MoA from traditional actin polymerisation inhibitors.

### Improving the potency of MMV291 conserves resistance to actin-1 and profilin in mutant parasites

We have previously reported the structure activity relationship (SAR) of MMV291 whereby the alpha-carbonyl *S*-methyl isomer was determined to be important for parasiticidal activity (Nguyen et al., 2021). Four of these analogues, *S-*W414, *S-*W936, *S-*W415 and *S-*W827 (Fig. S8) (previously referred to as *S-*18, *S-*20, *S-*22 and *S-*38) were selected to study the relationship of the chemical series targeting PfACT1 and PfPFN. These *S-*stereoisomers of the racemic MMV291 compound were tested on two clones from each chemically induced MMV291-resistant population in a 72-hour LDH growth assay. This revealed that the resistant clones maintained their resistance against the potent analogues of MMV291 (Fig. 3A-D). Interestingly, the degree of resistance differed depending on the parental population; population B clones (PFN(K124N)) were the least resistant, inducing a 10-fold increase in EC_50_ in the four analogues whilst the population C clones (ACT1(M356L)) exhibited the most resistance, increasing the EC_50_ 60 to 170-fold. This data indicated that since the ACT1(M356L) clones were consistently highly resistant to the MMV291 analogues, the MoA of this chemical series may be linked to PfACT1 function. biological experiments. Heat map indicates degree of resistance from 3D7 control lines, with yellow and red indicating the lowest and highest degree of resistance, respectively.

**Figure 3.**
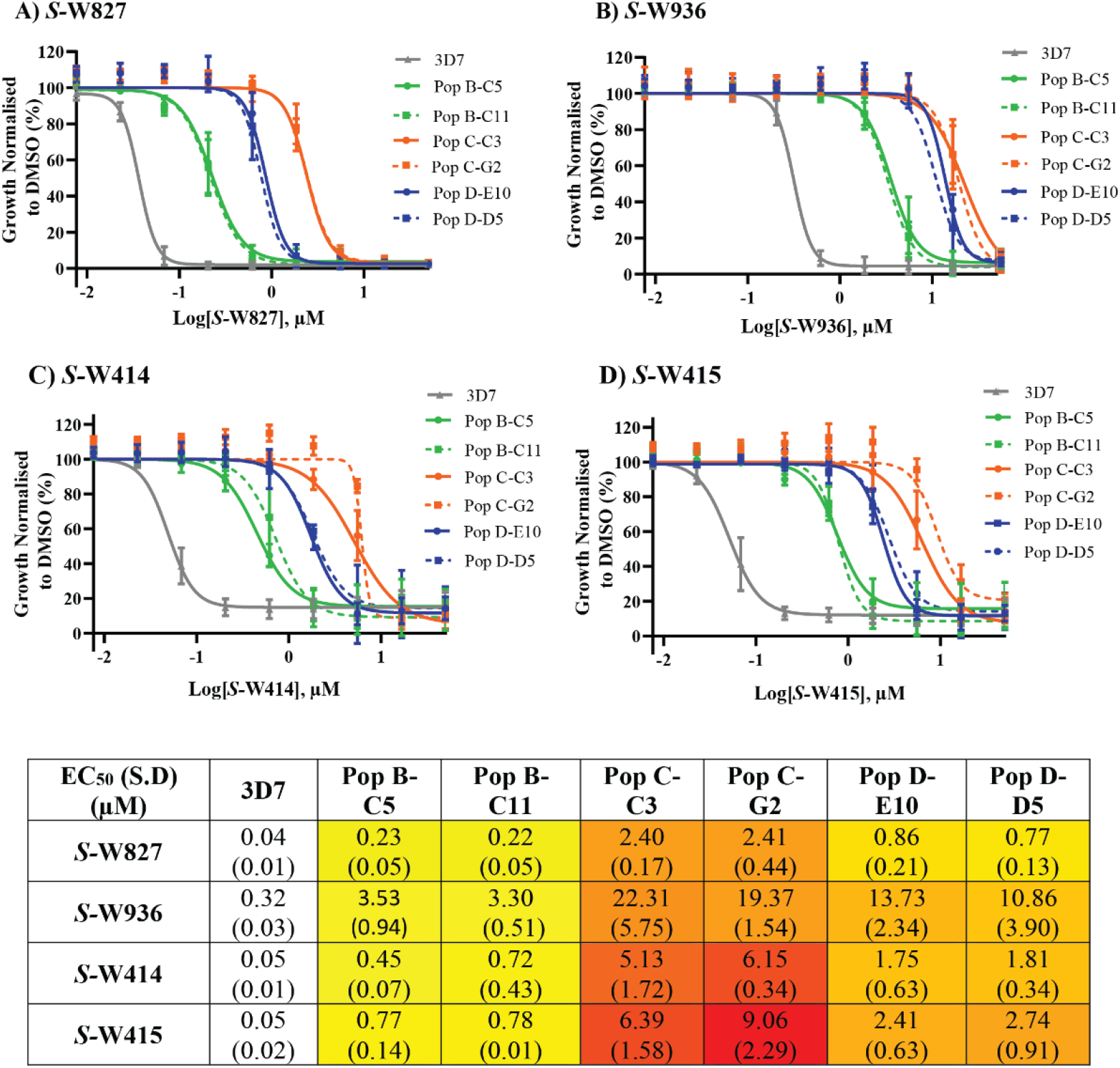
MMV291-resistant parasites demonstrate varying resistance to four analogues of MMV291. A) Two clones from three independently derived MMV291-resistant parasite lines were tested in 72-hour LDH growth assays. Varying degrees of resistance to *S-*W827 **(A)**, *S-*W936 **(B)**, *S-*W414 **(C)** and *S-*W415 **(D)** was observed, with Population C clones demonstrating the greatest resistance and Population B clones retaining the most sensitivity to the four molecules. Values were normalised to parasite growth in 0.1% DMSO, with error bars representing the mean of three biological replicates. Dose response curves were generated in GraphPad Prism using nonlinear regression to derive mean EC_50_ values which are stated in the table. S.D indicates the standard deviation calculated from EC_50_ values across three

### Actin polymerisation is affected by MMV291 *in vitro*

F-actin detection in apicomplexan parasites has been technically challenging because of the short length of the filaments produced (Pospich et al., 2017; Schmitz et al., 2010). Recently, this has been overcome with the expression of F-actin binding chromobodies in *Toxoplasma gondii* (Periz et al., 2017) that have also been adapted to *P. falciparum* (Stortz et al., 2019). The actin binding chromobodies consist of an F-actin nanobody fused to green fluorescent protein to allow microscopic detection of F-actin, which exists as a distinct punctate signal located at the apical tip of the merozoite. With actin polymerisation inhibitors, such as CytD, the punctate fluorescence dissipates into a uniform signal across the merozoite (Stortz et al., 2019). To ascertain if MMV291 could inhibit actin polymerisation in merozoites, we treated synchronised schizonts expressing the fluorescent nanobody with the parent MMV291 molecule and two analogues; *S-*W936, an active S-stereoisomer (EC_50_ of 0.2 µM), and *R-* W936, a less active R-stereoisomer of the former molecule (EC_50_ of 6.9 µM) (Fig. S8) at both 5× or 1× the growth EC_50_ (Fig. 4). *R-*W936 was also tested at multiples of *S-*W936 EC_50_ to allow a direct comparison between the compound’s activity and effect on actin polymerisation. Images of the egressed merozoites were captured and quantification of the punctate versus uniform F-actin signal was scored (Fig. 4A). This revealed that both concentrations of MMV291 and *S-*W936 tested, and high concentrations of less active isomer, *R-*W936, caused a similar reduction in merozoites expressing F-actin puncta to CytD treatment (P>0.05, Fig. 4B). In contrast, low concentrations of the less active isomer, *R-* W936, was significantly less effective at preventing merozoites from forming F-actin puncta than CytD (P<0.001, Fig. 4B). This result was notable as it provides the first direct link between the parasiticidal activity of MMV291, and its ability to inhibit F-actin formation in merozoites.

**Figure 4.**
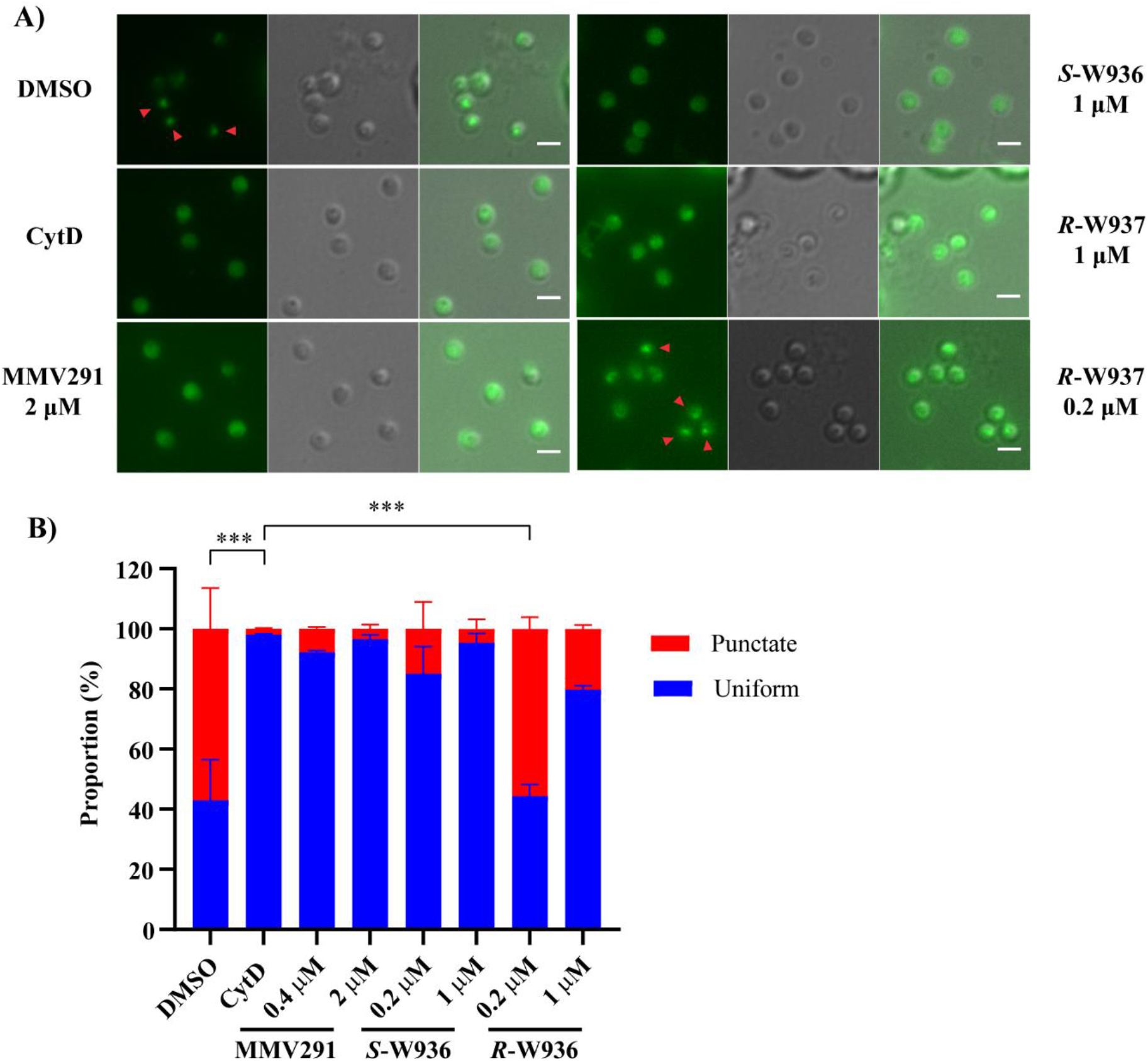
MMV291 treatment prevents F-actin formation in merozoitess. **A)** Synchronised schizonts from a *P. falciparum* parasite line expressing an F-actin-binding chromobody were incubated with DMSO, Cytochalasin D (CytD), MMV291 and analogues, *S-*W936 and *R-*W936, for 20 min at 37°C to allow merozoite egress. Merozoites were then imaged to detect either a normal punctate apical F-actin fluorescence signal or uniform signal, indicative of the inhibition of F-actin formation. Arrow heads depict punctate F-actin signal and scale bar indicates 2 μm. **B)** The proportion of merozoites with a punctate or uniform signal were scored with >550 merozoites counted for each treatment. Merozoites treated with the lower concentrations of the less active *R-*W936 had equal proportions of punctate and uniform fluorescence signals, like the DMSO control. In contrast, CytD, MMV291, and the active *S-*W936 compounds all greatly inhibited the formation of a punctate F-actin signal. Error bars represent the standard deviation of two biological replicates, each made up of three technical replicates from three individual counters. Statistical analysis was performed using a one-way ANOVA, comparing the mean of CytD punctate proportions with the mean of other treatments. *** indicates P<0.001, no bar indicates not significant. DMSO and CytD were used at concentrations of 0.1% and 1 µM, respectively.

Resistance to MMV291 arose due to mutations in both PfACT1 and PfPFN, suggesting the MMV291 series was interacting at the binding interface of the two proteins. To dissect the basis of this interaction, *in vitro* sedimentation assays with recombinant monomeric PfACT1 were carried out in the presence of compounds *S-*MMV291, *R-*MMV291, *S-*W936, *R*-W936, *S-*W414 and *S-*W827 and vehicle control, DMSO. In agreement with previous studies (Kumpula et al., 2019; Kumpula et al., 2017), 80% of PfACT1 could be sedimented into a pellet fraction by ultra-centrifugation in the absence of MMV291 analogues (Fig. 5A, Fig. S9A), whilst 15% of PfACT1 could be sedimented in the non-polymerizing (G-buffer) conditions (Fig. S9C, D). *S-*W936 was found to cause a small but significant reduction in the amount of PfACT1 in the pellet to 68% (P= 0.01, Fig. 5A). The remaining compounds had no statistically significant effect on PfACT1 sedimentation. These results indicate that the MMV291 analogues have either no or minimal impact on actin polymerisation *in vitro*.

**Figure 5.**
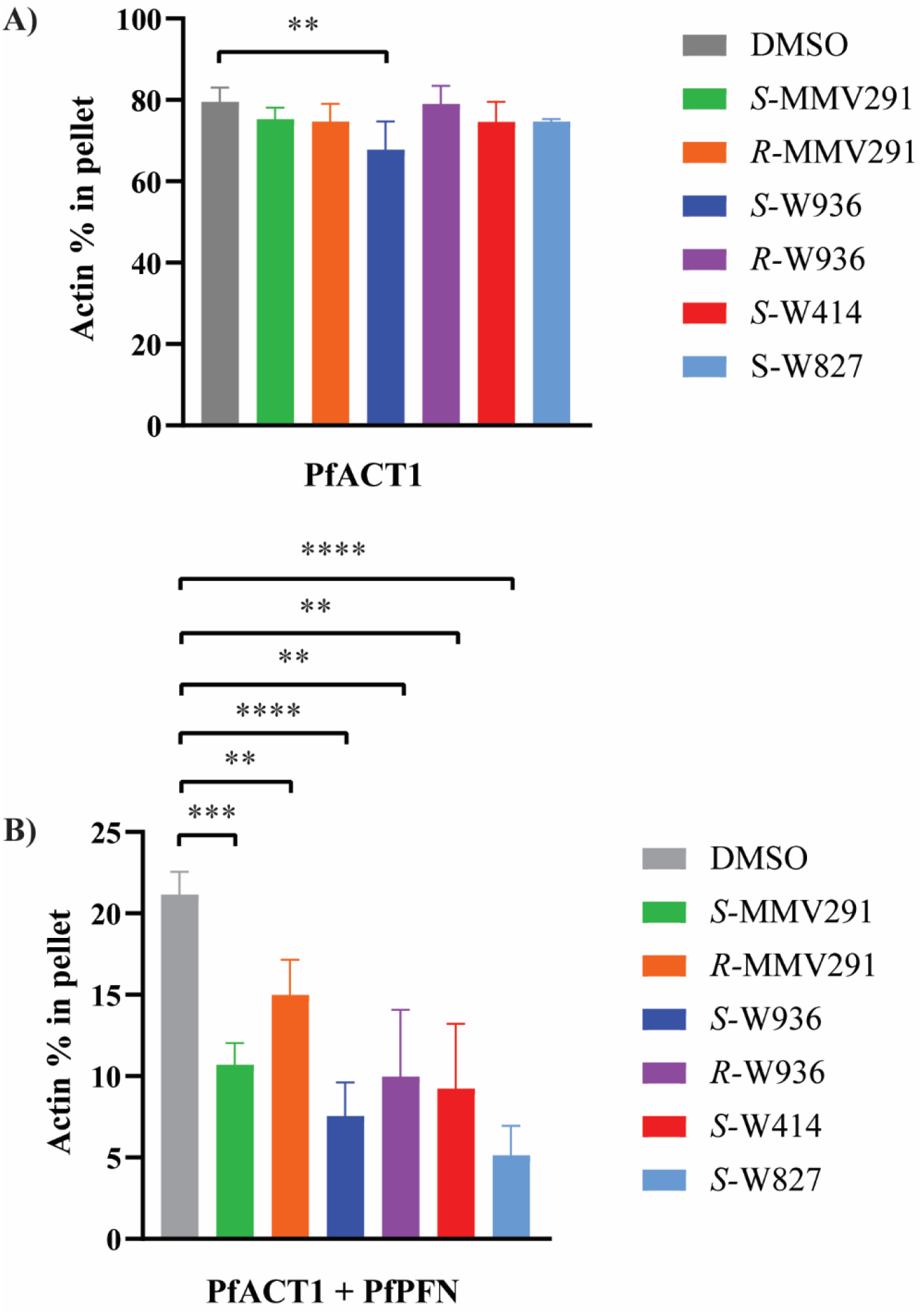
MMV291 analogues increase the actin-sequestering effect of profilin *in vitro*. PfACT1 (4 µM) under polymerizing conditions was quantified in the supernatant and pellet fractions in the presence of the MMV291 analogues (25 µM) or DMSO and upon addition of PfPFN (16 µM). **A)** In the absence of PfPFN, 80 ± 4% of PfACT1 sedimented to the pellet fraction with the vehicle DMSO treatment. *S-*W936 decreased the amount of actin in the pellet to 68 ± 7% whilst the remaining compounds had no significant effects on actin sedimentation. **B)** Upon addition of PfPFN, actin sedimentation decreased to 21 ± 1% with DMSO treatment. All MMV291 analogues, *S-*MMV291, *R-*MMV291, *S-*W936, *R-* W936, *S-*W414, and *S-*W827, decreased the amount of actin in the pellet further to 11 ± 1%; 15 ± 2%; 8 ± 2%; 10 ± 4%; 9 ± 4% and 5 ± 2%, respectively, indicating a stabilisation of the PfACT1-PfPFN complex. Results are plotted as mean ± standard deviation of the relative amounts of actin in the pellet fraction. The data are based on at least three independent assays each performed in triplicate. Statistical significances were determined using an unpaired two-tailed t-test, where ** P ≤ 0.01 and *** P ≤ 0.001, and **** ≤ 0.0001. No bar indicates not significant.

To address whether the MMV291 analogues affected the PfPFN-PfACT1 interaction, we included PfPFN in the sedimentation assays. As PfPFN sequesters G-actin, only 21% of PfACT1 remained in the polymerised pellet fraction following sedimentation (Fig. 5B, Fig. S9B). In the presence of the compounds, the amount of PfACT1 in the pellet decreased significantly to 7.5-15% with *S-*MMV291, *R-*MMV291, *S-*W936, *R-*W936 and *S-*W414 treatment (P< 0.01, Fig. 5B, Fig S9B). *S-*W827 exhibited the greatest affect by decreasing the PfACT1 to approximately 5% (P<0.0001). The magnitude of the effects observed from the different compounds on actin sedimentation was correlated with the EC_50_ values of the MMV291 analogues (Fig. S8) with the most potent inhibitors of parasite growth causing the greatest reduction in PfACT1 polymerisation. Notably, *R*-MMV291 had the smallest affect in agreeance with the weak parasite activity of this isomer compared to *S*-MMV291. In summary, these results indicate that the compounds may act through stabilising the PfACT1-PfPFN interaction, thus increasing the G-actin sequestering effect of PfPFN, while not having a significant effect on actin polymerization in the absence of PfPFN.

### MMV291 activity is specific for actin-1 dependent processes in the asexual stage of *P. falciparum* infection

F-actin is required for many processes across the lifecycle of *P. falciparum* including sporozoite gliding motility and hepatocyte invasion (Moreau et al., 2017; Münter et al., 2009). However, when sporozoites were treated with MMV291, both of these processes remained unaffected (Fig. S10). Similarly, despite the conserved sequences of actin-1 and profilin in *P. falciparum* and *T. gondii* (Fig. S11), MMV291 and its analogues also had little activity against tachyzoite invasion, unless the compounds were used at high concentrations relative to *P. falciparum* (500-1000 µM) (Fig. S12). Altogether this, combined with previous data showing MMV291 has little activity against gametocytes (Nguyen et al., 2021), indicates that this compound series has activity solely in the asexual stage of *P. falciparum* infection.

Next, we examined the effect of MMV291 on other F-actin dependent processes in the asexual stage. Conditional knockout of *actin-1* in *P. falciparum* have been shown to result in a defect in apicoplast segregation (Das et al., 2017). To evaluate the activity of MMV291 against apicoplast segregation, a parasite line that expresses a fluorescently tagged protein destined for trafficking to the apicoplast (ACP-GFP) was utilised (Waller et al., 2000).

Trophozoites were treated with 5 µM and 10 µM MMV291 (equating to 10×EC_50_ and 20×EC_50_) or the vehicle control before being imaged at schizont stages (Fig. 6Ai). The schizonts were scored to either have apicoplasts that were reticulated (an immature branched form), segregated (mature form), or clumped (abnormal) (Das et al., 2017; Waller et al., 2000). This revealed that the DMSO treatment resulted in a majority of normal apicoplast segregation with GFP labelling visualised as distinct punctate signals in daughter merozoites (Fig. 6Aii). In contrast, both concentrations of MMV291 induced a defect in apicoplast segregation whereby the 10 µM MMV291 resulted in significantly less segregated apicoplasts than the vehicle control (P=0.0003, Fig. 6Aii). This was visualised as distinct ‘clumps’, reminiscent of the phenotype shown previously in *actin-1* knockouts (Fig. 6A) (Das et al., 2017). Whilst the 5 µM concentration also displayed less segregation, this number was not significant (P=0.18, Fig. 6Aii). Altogether, this indicated that MMV291 induced a dose response effect on apicoplast segregation.

**Figure 6.**
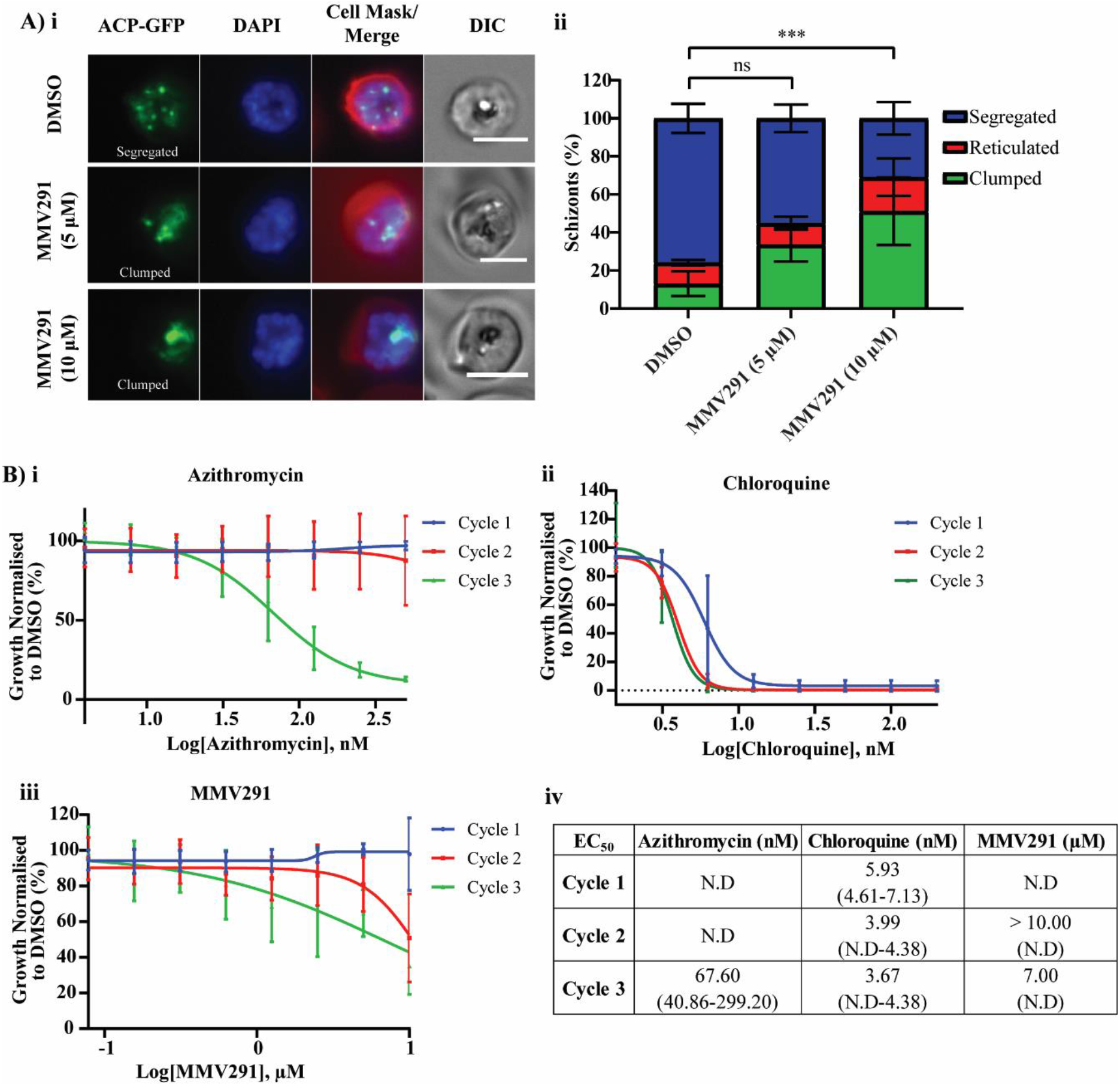
MMV291 disrupts actin-dependent apicoplast segregation and induces a partial delayed death phenotypeA) i. Representative panels from live cell imaging of apicoplast targeted acyl carrier protein (ACP) tagged with GFP, revealed that treatment of trophozoites for 24 hours with both 5 µM and 10 µM MMV291 (10 and 20×EC_50_) disrupted apicoplast segregation, resulting in an increase in abnormal apicoplast clumping at schizonts. Scale bar indicates 5 µm. **ii** These images were quantified by three independent blind scorers which showed a significant decrease in the normal segregation of apicoplasts between the 10 µM MMV291 and DMSO treatments, whilst the 5 µM MMV291 was not significant (ns). The number of cells analysed were 60, 48 and 47 for DMSO, 5 µM MMV291 and 10 µM MMV291, respectively which were captured over three biological replicates. The error bars represent the standard deviation of three independent blind scoring. Statistics were performed via a two-way ANOVA using GraphPad Prism between the DMSO segregated panel and the other treatments. *** indicates P<0.001. **B)** Early ring-stage exported nanoluciferase (Nluc) parasites were treated for 40 hours with a titration of azithromycin **(i)**, chloroquine **(ii)** or MMV291 **(iii)**, before compounds were washed out and the parasites allowed to grow until the second or third cycles with Nluc activity used to quantify parasite growth at each cycle. Unlike azithromycin, MMV291 did not display characteristics of a delayed death inhibitor but had partial reduction in parasite growth at the highest concentration used (10 µM) in the second and third cycles. In contrast, the fast-acting antimalarial chloroquine exhibited killing activity in the first cycle. Growth was normalised to that of parasites grown in 0.1% DMSO and EC_50_ values **(iv)** were derived from dose-response curves plotted from nonlinear regressions in GraphPad Prism with 95% confidence intervals of these values specified in brackets.

Defects in apicoplast inheritance for daughter merozoites induce a ‘delayed death phenotype’ whereby drugs targeting the apicoplast, such as the antibiotic azithromycin, exhibit no parasiticidal activity until the second cycle of growth after defective merozoites invade new RBCs and progress to trophozoites (Dahl and Rosenthal, 2007; Fichera and Roos, 1997; Kennedy et al., 2019). To investigate if MMV291 also produced a delayed death phenotype, highly synchronous ring-stage parasites expressing an exported nanoluciferase protein were treated with a titration of azithromycin, chloroquine or MMV291. After 40 hours and prior to merozoite invasion, the compounds were washed out and parasites allowed to grow for a further two cycles with nanoluciferase activity used as a marker for parasite growth. This demonstrated that azithromycin-treated parasites in cycle one elicited a dose-response decrease in parasite biomass in cycle three, producing an EC_50_ of 67.5 nM (Fig. 6Bi, iv), in contrast to chloroquine which demonstrated the profile of a fast-acting antimalarial (Fig. 6Bii). MMV291 displayed some intermediate delayed death activity at the maximum concentrations tested with cycle two and three producing EC_50_’s of >10 µM and 7 µM, respectively (Fig. 6Biii-iv). This was significantly higher than the compound’s overall EC_50_ of 0.5-0.9 µM in a 72-hour LDH assay, suggesting apicoplast segregation and subsequently delayed death is a secondary MoA of MMV291.

## Discussion

In this study, we sought to uncover the target and explore the MoA of a sulfonylpiperazine, MMV291, that acts to prevent merozoites from deforming and invading human RBCs. Resistance selection coupled with whole genome sequencing revealed three independent mutations in PfPFN and PfACT1 that did not impose a fitness cost on parasite growth *in vitro*. Furthermore, introducing these mutations into wildtype parasites mediated resistance to MMV291, indicating PfPFN and PfACT1 as putative targets of the compound. Interestingly, the three MMV291-resistant populations were observed to produce differing levels of resistance against the more potent MMV291 analogues, with parasites containing the PfACT1(M356L) mutation demonstrating the greatest resistance. This could indicate that MMV291 may interact with higher affinity to PfACT1 and thereby a conservative mutation may lead to reduced MMV291 binding, whilst still retaining the PfPFN-PfACT1 interaction. In contrast, the other two MMV291 PfPFN resistance mutations resulted in more radical amino acid changes and the fact that these mutants elicit similar overall parasite growth as the conservative PfACT1(M356L)-resistant parasites could indicate greater plasticity on the profilin side in PfPFN-PfACT1 binding.

We have previously reported that although MMV291-treated merozoites cannot deform and invade RBCs, the merozoites are still capable of irreversibly attaching to their target RBCs and can subsequently trigger echinocytosis (Dans et al., 2020). These characteristics are similar to those reported with CytD, although this naturally occurring compound acts upon the actin filament itself to prevent polymerisation (Shoji et al., 2012; Weiss et al., 2015). We previously noted that for RBCs with adherent MMV291-treated merozoites that the period of echinocytosis was greatly prolonged and that the adherent merozoites often produced pseudopodial extensions (Dans et al., 2020). Both effects have recently been observed in CytD-treated merozoites utilising a lattice light shield microscopy system (Geoghegan et al., 2021). Here, the authors described the membranous protrusions projecting from the parasite itself and extending into the interior of the RBC (Geoghegan et al., 2021). This CytD defect in merozoite invasion has also been reported as internal “whorls” which were visualised with antibody staining of the rhoptry bulb protein RAP1, indicating that although CytD blocks merozoite entry, rhoptry release was unaffected (Riglar et al., 2011). Disruption of RBC integrity due to the injection of merozoite rhoptry contents therefore appears to cause extended RBC echinocytosis unless the merozoite can enter the RBC and reseal the entry pore.

To further investigate the MMV291 series effect on actin polymerisation, *in vitro* actin sedimentation assays were carried out, revealing the compounds had no activity against PfACT1 polymerisation in the absence of PfPFN, apart from *S*-W936 that caused a slight reduction. However, all compounds tested significantly enhanced the ability of PfPFN to sequester actin monomers, with the greatest effects observed for the analogues which most potently inhibited parasite growth. It should be noted that although two of these analogues (*R-*MMV291 and *R-*W936) have low potency against the RBC stage of *P. falciparum* (EC_50_ >11 µM and 6.9 µM, respectively), sedimentation assays were carried out at 25 µM which would explain their activity in PfACT1 sequestration in the recombinant assay. This PfACT1 sequestration effect seen with the MMV291 analogues suggests that this compound series stabilises the interaction between PfACT1 and PfPFN, leading to decreased actin polymerisation. This could have a profound impact on the formation and turnover of F-actin required for invasion and other cellular functions. Indeed, the downstream effect of this stabilisation was observed in parasites expressing an F-actin chromobody whereby the MMV291 series was found to inhibit F-actin in merozoites in a manner that correlated with the parasiticidal activity of the compound. Altogether this forms the basis of our proposed model of the MoA of MMV291, whereby MMV291 increases the PfPFN sequestering effect of PfACT1, resulting in less PfACT1 turnover for the formation of the filaments, thereby functionally hindering the actomyosin motor and preventing merozoite invasion of RBCs (Figure 7).

**Figure 7.**
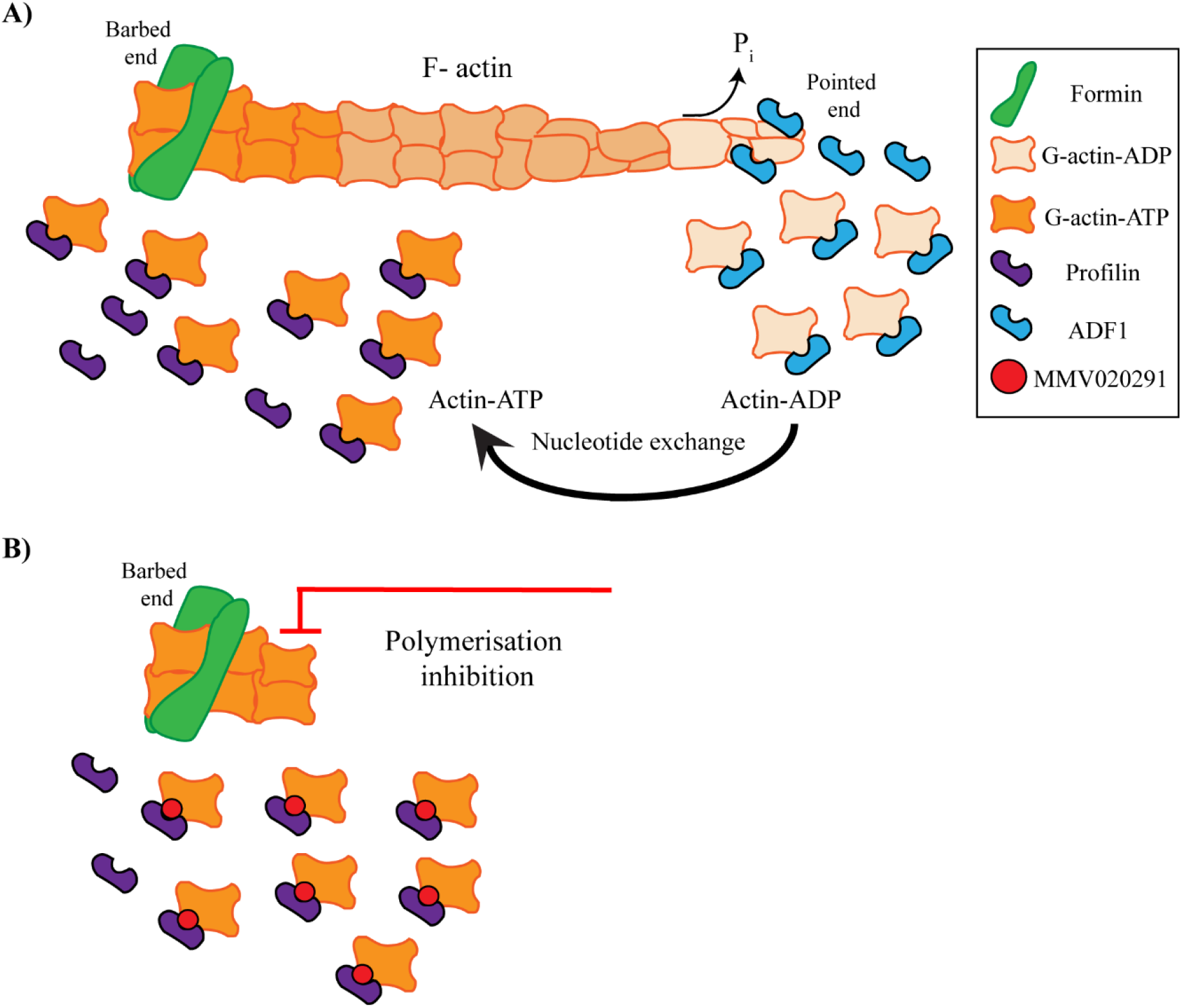
Proposed model for MMV291 interference in profilin-mediated filamentous actin polymerisation. **A)**Treadmilling model of profilin’s role in sequestering globular (G-actin) and stimulating the exchange of ADP for ATP before delivering the subunits to the barbed end of the growing filament. Here, formin initiates the polymerisation process to form filamentous actin (F-actin). Hydrolysis of the G-actin-ATP occurs at this end to produce G-actin-ADP and inorganic phosphate (P_i_), to stabilise the filament. The slow release of P_i_ at the pointed end induces filament instability and proteins such as actin depolymerising factor 1 (ADF1)bind to G-actin-ADP to aid in the release of the subunits, thereby severing the filaments.**B)** In the presence of profilin, MMV291 and its analogues inhibit the sedimentation of F-actin, suggesting the compounds stabilise the profilin-actin interaction to inhibit delivery G-actin to the barbed end. This inhibits F-actin polymerisation and thereby prevents merozoites from generating sufficient force required to invade RBCs.

This proposed MoA of PfPFN-ACT1 stabilisation within parasites contrasts a previously identified inhibitor of the profilin1/actin interaction in mammalian cells that was found to combat pathological retinal neovascularization (Gau et al., 2018; Gau et al., 2020). Here, the authors confirmed the competitive activity of the compound by demonstrating its inhibition of actin polymerisation in the presence of profilin1 (Gau et al., 2018), highlighting the druggable potential of this protein-protein interaction. It is thought that apicomplexan profilin may have originated from an evolution fusion of two ancestral genes (Bhargav et al., 2015) and therefore the PfPFN-ACT1 interaction may provide the basis of a selective drug target not found in their mammalian counterparts. This was reinforced by the lack of activity of MMV291 against HepG2 cells (Nguyen et al., 2021) or merozoite invasion into RBCs pre-treated with MMV291.

Despite the phenotype of MMV291-treated merozoites phenocopying CytD, the MoA of MMV291 interference in actin polymerisation is more reminiscent of the latrunculins. These naturally occurring compounds prevent actin turnover through binding to G-actin subunits (Morton et al., 2000; Varghese et al., 2020). Whilst targeting both G-actin and PFN-ACT1 result in a similar phenotype of reduced filament formation, compounds directed at the PfPFN-ACT1 interaction may have more success due to greater selectivity. Crystallography studies solving the binding site of the MMV291 series in relation to the PfPFN-ACT1 interaction would be worthwhile undertaking in order to gain a greater understanding of the druggable potential of these essential parasite proteins.

Whilst crucial for merozoite invasion, PfPFN-PfACT1 may not be required for other F-actin dependent processes such as gametocytogenesis and apicoplast segregation (Das et al., 2017; Das et al., 2021; Hliscs et al., 2015). The MMV291 series have previously shown little activity against gametocytes (Duffy et al., 2017; Nguyen et al., 2021) and whilst we did observe some activity against apicoplast segregation with MMV291 treatment, this parasiticidal activity occurred in much greater concentrations than observed within a standard 72-hour growth assay. This implicates apicoplast segregation as a secondary MoA of MMV291 and perhaps other G-actin sequestering-binding monomers, such as actin-depolymerisation factor 1 (ADF1) (Schüler et al., 2005; Wong et al., 2011), could be the predominant PfACT1 sequesters that are utilised by parasites for these F-actin dependent processes. Additionally, the requirements for PfACT1 sequestering and subsequent turnover of F-actin may vary dependent on the process at hand. This can be realised by the varying speeds in motility in different stages of parasites; ookinetes move at 5 µm/min (Kan et al., 2014; Vlachou et al., 2006; Vlachou et al., 2004), merozoites are the next fastest at 36 µm/min (Yahata et al., 2021), whilst the fastest are sporozoites which can reach speeds of 60-120 µm/min (Hopp et al., 2015; Ripp et al., 2021).

It was somewhat unexpected that MMV291 did not reduce sporozoite motility since sporozoites have been shown to be highly sensitive to mutations in profilin (Moreau et al., 2017; Moreau et al., 2021). However, these mutations were located in an acidic loop and a conserved β-hairpin domain, which led to the disruption or weakening of the PFN-ACT1 complex and thereby implicating this interaction as an essential requirement for fast-gliding motility (Moreau et al., 2017). It is therefore possible that stabilising the PFN-ACT1 interaction is not detrimental to actin polymerisation within sporozoites. Furthermore, we showed that hepatocyte invasion of sporozoites were unaffected by MMV291 treatment. This may be due to the different requirements for the PFN-ACT1 interaction to aid in actin polymerisation and subsequent G-actin turnover to invade these host cells with varying membrane tensions and elasticity. Altogether, this indicates that stabilising the PfACT1-PfPFN interaction appears to specifically inhibit *P. falciparum* invasion of RBCs.

This trend of specificity for merozoite invasion of RBCs was extended to *T. gondii* where tachyzoites also displayed limited sensitivity to the MMV291 series. Here, high concentrations of compounds were required to elicit a reduction host cell invasion. Despite TgPFN being essential for host cell invasion (Plattner et al., 2008) and that TgPFN and PfPFN proteins are somewhat conserved (Moreau et al., 2017), a key difference between these parasites is that TgPFN inhibits the conversion of ADP-ATP on G-actin, thereby inhibiting F-actin polymerisation (Kucera et al., 2010; Skillman et al., 2012). This has not been observed with PfPFN (Das et al., 2021; Moreau et al., 2017). This highlights the diverged nature of profilin within apicomplexan parasites and, along with differences in host cells, may explain the disparity in activity of the series between *P. falciparum* and *T. gondii*.

Within the *Plasmodium* spp, profilin is highly conserved (Kursula et al., 2008). MMV291 has previously been shown to possess activity against *Plasmodium knowlesi*, albeit with less potency than *P. falciparum* (Nguyen et al., 2021), indicating that there may be a conserved PFN-ACT1 mechanism across *Plasmodium* spp. that is required for invasion of RBCs. Indeed, the resistant mutation locations are conserved in *P. knowlesi* profilin (PkPFN(K125), PkPFN(N155)) but further work as to whether this parasiticidal activity is linked to invasion defects in *P. knowlesi*, and if it extends to other *Plasmodium* spp., is required.

In summary, this investigation identified the first specific inhibitor of *P. falciparum* actin polymerisation in which its MoA could be linked through interference with PfPFN/ACT1 dynamics. Further investigations into the binding locations of MMV291 within its target proteins are warranted as this may shed more light onto the exact mechanism(s) by which the compound acts to inhibit merozoite invasion. The antimalarial development of this compound series is currently hampered by the high clearance of MMV291 and its analogues in liver microsomes (Nguyen et al., 2021). Additional medicinal chemistry work is therefore required to address the metabolic instability of this series before it can progress further towards a future antimalarial. Nonetheless, the MMV291 series could serve as a useful tool to study the complex regulation of actin polymerisation in the malaria parasite. This, in turn, could provide a starting point for future development of novel scaffolds against profilin-mediated F-actin polymerisation.

## Acknowledgements

We acknowledge the Australian Red Cross Blood Bank for the provision of human blood. We thank D. Marapana for the p1.2 CRISPR plasmid. We thank S. Tan for the PbCSP antibody and J. Boddey for the PfCSP antibody.

## Materials/Methods

### *Plasmodium falciparum in vitro* culturing and parasite lines

*Plasmodium falciparum* parasites were cultured as previously reported (Trager and Jensen, 1976) in human O type RBCs (Australian Red Cross Blood Bank) at 4% haematocrit in RPMI-HEPES supplemented with 10% v/v heat-inactivated human serum (Australian Red Cross) or albumax (Gibco). Unless specified, all assays using *P. falciparum* were conducted on laboratory wild-type strain 3D7 parasites. An exported Nanoluciferase (Nluc) parasite line (ex-Nluc) (Azevedo et al., 2014) was used for Nluc-based assays and episomally maintained using 2.5 nM WR99210 (Jacobus Pharmaceutical Company). For apicoplast segregation assays, a parasite line expressing a green fluorescent protein-tagged acyl carrier protein (ACP-GFP) (Waller et al., 2000) was used, and this was maintained episomally in parasites by addition of 0.1 µM pyrimethamine (Sigma Aldrich). A chromobody-emerald fluorescent protein expressing *P. falciparum* parasite line was used for actin-chromobody experiments (Stortz et al., 2019).

### Generation of *P. falciparum* sporozoites

Gelatine selected *P. falciparum* parasites (NF54e strain, Walter Reed National Military Medical Center, Bethesda) were utilised for gametocyte generation as previously described in (Saliba and Jacobs-Lorena, 2013) with daily media changes. Gametocytes from these cultures were diluted to 0.1% and 0.3% gametocytaemia and fed to *Anopheles stephensi* mosquitoes on artificial membrane feeders. *A. stephensi* mosquitoes (STE2, MRA-128, from BEI Resources) were reared in an Australian Biosecurity (Department of Agriculture and Water Resources) approved insectary. The conditions were maintained at 27°C and 75–80% humidity with a 12-hour light and dark photo-period in filtered drinking water (Frantelle beverages, Australia) and fed with Sera vipan baby fish food (Sera). The larvae were bred in plastic food trays (cat M612-W P.O.S.M Pty Ltd, Australia) containing 300 larvae, each with regular water changes every three days. On ecloding, the adult mosquitoes were transferred to aluminium cages (cat 1450A, BioQuip Products, Inc.2321 Gladwick St.Rancho Dominguez, CA 90220) and kept in a secure incubator (Conviron), in the insectary at the same temperature and humidity and maintained on 10% sucrose. Fed mosquitoes were kept at 27 °C in a humidified chamber (Goodman et al., 2016). Salivary glands of infected mosquitoes (day 21 post-infection) were isolated by dissection and parasites placed into RPMI-1640 media.

### Generation of *Plasmodium berghei* sporozoites

*Plasmodium berghei* ANKA wild-type Cl15cy1 (BEI Resources, NIAID, NIH: MRA-871, contributed by Chris J. Janse and Andrew P. Waters) was used for sporozoite motility assays.

*P. berghei* ANKA reporter parasite lines for the *in vitro* liver stage drug assay PbGFP-Luc con (676m1cl1) RMgm-29 (http://www.pberghei.eu/index.php?rmgm=29) was provided by Andy Waters (University of Glasgow, Glasgow, Scotland) (Annoura et al., 2013).

Animals used for the generation of the sporozoites were 4-5 week old male Swiss Webster mice and were purchased from the Monash Animal Services (Melbourne, Victoria, Australia) and housed at 22-25°C on a 12 hour light/dark cycle at the School of Biosciences, The University of Melbourne, Australia. All animal experiments were in accordance with the Prevention of Cruelty to Animals Act 1986, the Prevention of Cruelty to Animals Regulations 2008 and National Health and Medical Research Council (2013) Australian code for the care and use of animals for scientific purposes. These experiments were reviewed and permitted by the Melbourne University Animal Ethics Committee (2015123).

Infections of naïve Swiss mice were carried out by intraperitoneal (IP) inoculation obtained from a donor mouse between first and fourth passages from cryopreserved stock. Parasitemia was monitored by Giemsa smear and exflagellation quantified three days post infection. 1 μL of tail prick blood was mixed with 100 µL of exflagellation media (RPMI-1640 [Invitrogen] supplemented with 10% v/v foetal bovine serum (FBS), pH 8.4), incubated for 15 min at 20°C, and exflagellation events per 1×10^4^ RBCs were counted. *Anopheles stephensi* mosquitoes were allowed to feed on anaesthetised mice once the exflagellation rate was assessed between about 12-15 exflagellation events per 1×10^4^ RBCs. Salivary glands of infected mosquitoes (days 17–24 post-infection) were isolated by dissection and parasites placed into RPMI-1640 media.

### Compounds

Stock concentrations of Azithromycin (20 µg/µL, AK-Scientific), CytD (2 mg/mL, Sigma Aldrich), Jasplakinolide (1 mM, Sigma Aldrich), ML10 (10 mM, Lifearc) were made up in DMSO (Sigma Aldrich). Chloroquine (10 mM, Sigma Aldrich) and Heparin (100 mg/mL, Sigma Aldrich) were dissolved in H_2_O and RPMI, respectively. MMV291, *S-*MMV291, *R-* MMV291, *S-*W936, *R-*W936, *S-*W414, *S-*W415 and *S-*W827 (Walter and Eliza Hall Institute) were dissolved in DMSO to a 10 mM stock solution.

### Resistance selection

Resistant parasites were generated as previously described (Gilson et al., 2019) by exposing a clonal population of 10^8^ 3D7 wildtype (WT) parasites to 10 μM MMV291 (∼10×10EC_50_).

The parasites were incubated with the compounds until the drug treated parasites began to die off, with the drug replenished daily. The compounds were then removed until healthy parasite replication was observed via Giemsa-stained thin blood smear, upon which compound treatment was resumed. Altogether, the compounds were cycled on and off for three cycles until three populations of MMV291 were observed to be resistant to the compounds via a growth assay. The resistant lines were cloned out by limiting dilution prior to genomic DNA (gDNA) extraction and their EC_50_ for growth was evaluated following a 72-hour treatment to ensure the resistance phenotype was stable.

### Whole genome sequencing and genome reconstruction

Late-stage parasites from the original 3D7 clonal line and MMV291-resistant clones were harvested via saponin lysis (0.05%, Kodak). A DNeasy Blood and Tissue kit (Qiagen) was then used to extract gDNA from the saponin-lysed pellets following the kit protocol with the exception that additional centrifugation steps were performed to remove hemozoin prior to passing lysates through the DNA binding columns.

Concentration of extracted DNA was evaluated by Qubit Fluorometer (Invitrogen Life Technologies). Average length of the DNA sample was then assessed using Tapestation (Agilent Technologies). Based on concentration and average length of DNA sample, 0.2 pmol DNA was carried forward to prepare sequencing library using 1D genomic sequencing kit SQK-LSK109 and Native barcodes (EXP-NBD104, EXPNBD114) as per manufacturer’s instructions (Oxford Nanopore Technologies, UK). For maximum sequencing output, each sequencing run comprised of 3-5 samples labelled with distinct Oxford Nanopore native barcodes. A 48-hour sequencing run was performed on a MinION platform with MIN106D Flow cells and MinIT (Software 18.09.1) to generate live fastq files.

After sequencing, fastq files were subjected to demultiplexing and adapter trimming was subsequently performed using Porechop (V0.2.3_seqan2.1.1). Read alignment against the *P. falciparum* 3D7 reference genome was performed using minimap2 (V2.17) default parameters. Alignment files (sam format) were processed with samtools utilities (V1.9) to generate sorted bam files. Genotype likelihoods were then computed using bcftools mpileup (V1.9), configured to skip indels and consider only bases with quality 7 or above. Variant calling was then performed using bcftools multiallelic-caller (V1.9) to generate haploid SNP candidates for each sequenced isolate. Candidate SNPs in blacklisted genomic regions (Miles et al., 2016) (that is, subtelomeric, centromeric or internal hypervariable regions) were removed using bedtools subtract (V2.29.2) to retain SNPs in the core genome only. Quality filtration of SNP calls, enforcing a minimum depth of coverage of 10X and requiring at least 70% of reads to support the called allele (Razook et al., 2020), was performed using custom awk script.

Filtered candidate SNPs for each isolate were then imported into R statistical software (V3.6.1) and overlaps between the 3D7 WT and resistant isolates were examined. To account for differences between the 3D7 reference isolate and our independently cultured 3D7 WT isolate, SNPs present in the 3D7 WT isolate were removed. Functional annotation of non-WT candidate SNPs was performed using the VariantAnnotation package (V1.32.0), retaining only non-synonymous SNPs. To identify causal resistance variants, biological annotations, including gene ontology terms and expression profiles, were collated for the final set of candidate SNPs using in-house software (PlasmoCavalier).

### Data Availability

Genomic sequencing data is available from European Nucleotide Archive; accession number PRJEB55647.

### Molecular biology and transfection of *P. falciparum*

To introduce the *profilin* resistant mutations, the donor plasmid was designed whereby the coding sequence of *profilin* (PF3D7_0932200) was recodonised without introns to the bias of *Saccharomyces cerevisiae* and synthesised as gBlock fragments (Integrated DNA technologies) for both the WT and N154Y sequences. The cloning strategy involved amplifying the first 684 bp of the 5’ native sequence of *profilin* using gDNA to form homology region 1A (primers listed in Table S1, a/b) and the 3’ recodonised gBlock fragment comprising of the last 292 bp of either the WT or N154Y *profilin* to form homology region 1B (primers listed in Table S1, c/d). The two products were sewn together using primers listed in Table S1 (a/d) to reconstitute the full coding sequence of *profilin* (HR1). The 3’ flank of the native *profilin* sequence was amplified using primers stated in Table S1 (e/f) to create the 3’ homology region 2 (HR2). To introduce the K124N mutation, primers outlined in Table S1 (g/h) were used to generate a HR1(K124N) using the WT sequence as a template. Both HR1(WT/N154Y/K124N) and HR2 were digested with *Bgl*II/*Spe*I or *EcoR*I/*Kas*I, respectively and inserted into the parasite vector p1.2 (Marapana et al., 2018).

To introduce the *actin-1* resistant mutation, *actin-1 (PF3D7_1246200)* was recodonised to a *Spodoptera frugiperda* bias and synthesised as a gBlock fragment (Integrated DNA technologies). To create the full length HR1, the first 517 bp of the 5’ end of *actin-1* (HR1A) and 614 bp of the 3’ recodonised gBlock (HR1B) were amplified using gDNA and the gBlock fragment as templates, respectively (primers in Table S1 l/m, n/o). The two fragments were then sewn together using primers listed in Table S1 (l/o) to reconstitute the coding sequence. The M356L mutation was introduced using overlapping primers as stated in Table S1(r/s). The HR2 sequence of *actin-1* was amplified from gDNA using the primers listed in Table S1 (p/q). An internal *Bgl*II site in the HR2 of the native *actin-1* sequence was repaired using overlapping primers in Table S1 (t/u). Both HR1(WT/M356L) and HR2 were introduced into parasite vector p1.2 as outlined above.

The guide RNA (gRNA) sequences were designed with a protospacer adjacent motif (PAM) site using the online program https://chopchop.cbu.uib.no/ (Labun et al., 2019) (sequences in Table S1). The donor constructs were linearized and transfected into WT 3D7 ring-stage parasites along with recombinant Cas9 enzyme and annealed gRNA and tracrRNA (Integrated DNA technologies) as described in (McHugh et al., 2021). Chromosomal integration of the construct, which includes the human dihydrofolate resistance gene (hDHFR), was selected for with 2.5 nM WR99210. Once viable parasites were obtained, gDNA was extracted and integration PCRs were performed with expected products for modified and parental loci (primers listed in Table S1). These PCR products were sequenced (Micromon Sanger sequencing) to confirm presence of resistant alleles.

### Delayed death assay

The assay was adapted from a previously described method (Kennedy et al., 2019) and utilised ex-Nluc parasites that had been tightly synchronised using 25 nM ML10. Ring-staged parasites (0-4 hours post invasion) were diluted in 96 well U-bottom plates to 1%, 0.1% and 0.01% parasitemia for cycle one, cycle two and cycle three, respectively, each with three technical replicates. The parasites were then distributed into serially diluted MMV291, azithromycin and chloroquine from 10 µM, 0.5 µM and 0.2 µM respectively, in a two-step dilution. Parasites were incubated with the compounds for approximately 40 hours until they reached the late-trophozoite to early-schizogony stage and cycle one plates were frozen. The remaining plates were washed three times before pellets were resuspended to a final volume of 100 μL. The plates were then incubated at 37°C for a further 48 hours before cycle two plates were frozen. Cycle 3 plates were grown for a further 48 hours before also being frozen. At the completion of the assay, plates were thawed and the Nluc signal was quantified by adding 5 µL of whole iRBCs to 45 µL of 1 x Nanoglo Lysis buffer with 1:1000 NanoGlo substrate (Promega) in a white luminometer 96-well plate. A CLARIOstar luminometer (BMG Labtech) was used to measure relative light units (RLU) and growth was normalised to 0.1% DMSO.

### Pre-treatment RBCs assay

Fresh RBCs (2 µL of 50% haematocrit) were added to either 100 µL of 0.1% DMSO, 10 µM MMV291, 100 µg/mL heparin or 0.0025% glutaraldehyde (ProSciTech) in a 96-well U-bottom plate and incubated for 30 min at 37°C, with shaking at 400 rpm. RBCs were then washed three times and resuspended in a final volume of 50 µL. Invasion assays were then carried out as previously described in (Boyle et al., 2010; Dans et al., 2020; Wilson et al., 2015; Wilson et al., 2013) whereby purified merozoites expressing an ex-Nluc were then added to pre-treated RBCs and naïve RBCs (i.e., RBCs that had not been pre-treated with compounds). Invasion efficiency was then evaluated as described in (Dans et al., 2020).

### Lactate dehydrogenase (LDH) malstat growth assay

These were performed as previously described in (Makler and Hinrichs, 1993). Synchronous ring-staged parasites were diluted to 0.3% parasitemia and added to serial dilutions of compounds in a 96 well U-bottom plate, with a final haematocrit and volume of 2% and 100 μL, respectively. Plates were incubated for 72 hours at 37°C and were then frozen at -80°C until assayed. After thawing, 35 µL of parasite culture was added to 75 µL of Malstat reagent in a 96 well flat-bottom plate and incubated in the dark for 30-60 min until colour change occurred. Absorbance (650 nm) was measured on a Multiskan Go plate reader Thermo Scientific), using Skan IT software 3.2. Growth was then expressed as a percentage of vehicle control and dose-response curves plotted using GraphPad Prism 8.4.0.

For the multicycle growth assays, the parasitemia of ring-stage MMV291 resistant clones, E10, B11 and C3, and 3D7 parasites were counted and adjusted to 0.3% parasitemia with 2% haematocrit. Over 10 cell cycles, samples were taken at each cycle and frozen until completion of the assay. To ensure overgrowth of parasites did not occur, at each cycle, parasites were diluted 1 in 8 which was accounted for in the analysis. Growth was measured by a LDH growth assay as outlined above.

### Actin-binding chromobody assay

A *P. falciparum* 3D7 parasite strain expressing a chromobody-emerald fluorescent protein (Stortz et al., 2019) was synchronised using Percoll (Sigma Aldrich) purification and sorbitol lysis and grown for 45 hours to schizont stages. 2 nM of Compound 2 was administered to schizonts and incubated for 3 hours at 37°C. The inhibitor was then washed out and schizonts were returned to pre-warmed complete RPMI media containing either MMV291, *S*-936, *R*-936, CytD or DMSO and added into a microscope chamber. Here, schizonts were incubated at 37°C for 20 min to allow merozoite egress before live imaging of newly egressed merozoites were conducted. Quantification of images was conducted by three independent scorers.

### Apicoplast segregation assay

ACP-GFP parasites (Waller et al., 2000) were treated at trophozoite-stage with 5 or 10 µM MMV291 or the vehicle control for 24 hours until they reached schizogony stage. Prior to imaging, they were incubated with 5 µg/mL CellMask DeepRed (Thermo Fisher Scientific) and 0.3 μM of 4′,6-diamidino-2-phenylindole (DAPI) in RPMI with decreased albumax (0.13%) in complete RPMI for 30 min. The cells were then washed three times in complete RPMI, mounted and imaged on a Zeiss Cell Observer widefield fluorescent microscope. Apicoplasts were scored by three independent blinded scorers as fully segregated, reticulated (branched) or clumped (not segregated). Statistical tests were performed in GraphPad Prism 8.4.0 using a two-way ANOVA with multiple comparisons between each treatment group.

### Sporozoite motility assay

Eight-well chamber slides (PEZGS0816 Millipore Millicell EZ SLIDE) were coated with a monoclonal antibody (mAb) specific for the repeat region of the circumsporozoite protein 3D11 mouse anti-PbCSP (RRID:AB_2650479) (Yoshida et al., 1980) at a 1 in 1000 dilution in phosphate buffered saline (PBS) for 30 min at 37°. 8,000-10,000 sporozoites were incubated in 100 µL of RPMI 1640-HEPES Glutamax (Invitrogen) supplemented with 10% heat-inactivated FBS (Invitrogen), with the appropriate drug concentrations and solvent controls for 30 min at 22°C. The sporozoites were then seeded into a coated well per treatment and allowed to glide for 45 min at 37°C in an 5% CO_2_ incubator. Experiments were stopped by removing the supernatant and fixing with 4% v/v paraformaldehyde in 1× PBS at 37°C for 15-20 min. Primary antibody of PbCSP (courtesy of S. Tan) or PfCSP (courtesy of J. Boddey) (1/1000 dilution in 3% bovine serum albumin (BSA) in 1×PBS) was applied for 45 mins followed by a secondary antibody goat anti-mouse IgG, Alexa fluor R 488; (Invitrogen) (1/1000 dilution in 3% BSA in 1×PBS). After each of these stages, the wells were washed gently with 1x PBS. Finally, Hoechst 33342 (Sigma Aldrich) in 1× dPBS was added to each of the wells at a final concentration of 5 μg/mL, incubated for 5 min, washed with dH_2_0 and air dried. The number of sporozoites with and without trails was counted under a fluorescent microscope (Olympus CKX41) after 10 µL of DAKO (Sigma Aldrich) and a coverslip were applied.

### *In vitro* liver stage assay

This was performed as described in (Annoura et al., 2013) but with the following variations. *In vitro* human liver HCO4 cells (ATCC) were seeded at 1×10^5^ cells/mL, 200 µL in each well in a 96 well Nunc™ Edge™ 96-Well, Nunclon Delta-Treated, Flat-Bottom Microplate (Thermo) and grown for 24 hours in Advanced MEM (Invitrogen), 10% (vol/vol) FBS (Invitrogen), and 1% (vol/vol) antibiotic–antimycotic (Invitrogen) in a standard tissue culture incubator (37 °C, 5% CO_2_). 20,000 sporozoites from freshly dissected infected mosquitoes were added per well. After 52 hours, the supernatant was removed gently, the wells washed with 1x PBS and after the removal of the PBS, the plate was placed in the -80°C freezer for at least 30 min. To perform the luciferase assay, the plate was removed from the -80°C and 20 µL of 1× cell culture lysis reagent (CCLR Promega, cat. no. E1531) was added to the frozen plate. The plate was shaken at room temperature for 15-20 min. This lysate was transferred to Nunc™ MicroWell™ 96-Well, Nunclon Delta-Treated, Flat-Bottom Microplate (cat: 236105 Thermo Scientific) 20 µL of the Luciferase assay substrate solution (Luciferase Assay System Kit Promega, cat. no. E1500) was added into each of the wells of the lysed samples. RLU for each sample was then measured via a micro plate reader (EnSpire Perkin Elmer).

### *Toxoplasma gondii* invasion assays

Freshly egressed Nluc expressing parasites were harvested and passed through a 25-gauge needle three times to liberate from host cells. Parasites were counted and then preincubated for 20 min with different concentration of the compounds, ranging from 1000 µM to 8 µM diluted in DMEM supplemented with 5% FBS. Parasites were then transferred into 96-well plates containing human foreskin fibroblasts in triplicate and centrifuged at 290 g for 5 min at room temperature. Plates were then transferred to an incubator and the parasites were allowed to invade for 1 hour at 37°C in the presence of 10% CO_2_. Wells were then washed with DMEM four times to remove any non-invaded parasites. Plates were then placed at 37°C with 10% CO_2_ for 24 hours in 200 µL DMEM supplemented with 5% FBS. Invasion media was then removed and the host cells containing the Nluc expressing parasites were then lysed using Promega Nano-Glo luciferase assay kit and the light units quantified on a Millennium Science plate reader.

### Recombinant protein production and actin sedimentation assays

Wild-type PfPFN cloned into pET28a(+)-TEV using NdeI/BamHI cloning site was ordered from GenScript (Leiden, Netherlands), expressed *in E. coli* BL21(DE3) cells, and purified using standard protocols as described in (Moreau et al., 2021). As an exception, the purification tag was cleaved with TEV during dialysis. PfACT1 was produced in *Spodoptera frugiperda* Sf21 cells (Invitrogen) as described in (Ignatev et al., 2012), with a few changes in the protocol. When infecting the cells, 13.5 µL of high-titer virus was used per 10^6^ cells, the cells were harvested 4 days after infection and used right away for protein purification as described in (Moreau et al., 2017).

Sedimentation of 4 µM PfACT1 1 in 10 mM HEPES pH 7.5, 0.2 mM CaCl_2_, 0.5 mM ATP, 0.5 mM TCEP and 2.5% DMSO was studied alone and in the presence of 25 µM MMV291 analogues, or in the presence of 16 µM PfPFN without and with 25 µM MMV291 analogues. Actin polymerization was induced by adding polymerizing buffer to final concentrations of 50 mM KCl, 4 mM MgCl_2_ and 1 mM EGTA. For control purposes, PfACT1 samples without polymerizing buffer were included to the assay. Total sample volume was 150 µL. Samples were polymerized overnight (∼16 hours) at room temperature (∼22°C), 100 µL of each sample (in triplicate) were centrifuged for 1 hour at 20°C using 100,000 rpm and TLA-100 rotor (Beckman Coulter, CA, USA). The resulting supernatants and pellets were separated, the supernatants were mixed with 25 µL of 5x SDS-PAGE sample buffer (250 mM Tris-HCl pH 6.8, 10% SDS, 50% glycerol, 0.02% Bromophenol Blue and 1.43 M β-mercaptoethanol), and the pellets were suspended in 125 µL of 10 mM HEPES pH 7.5, 0.2 mM CaCl_2_, 0.5 mM ATP, 0.5 mM TCEP supplemented with 1× SDS-PAGE sample buffer. Samples were incubated 5 min at 95°C, and then 10 µL of each sample was analysed on 4–20% Mini-PROTEAN TGX gel (Bio-Rad Laboratories, CA, USA). The protein bands were visualized with PageBlue stain (Thermo Scientific, MA, USA). Gels were imaged using the ChemiDoc XR S+ system (Bio-Rad) and protein band intensities were determined with the ImageJ 1.52D software (Schneider et al., 2012). For each supernatant and pellet pair, the relative amounts of PfACT1 in pellets were presented as percentages of the total intensity of PfACT1 which was set to 100%.

