## Supplementary Figures and Table for "The sulfonylpiperazine MMV020291 prevents red blood cell invasion by the malaria parasite *Plasmodium falciparum* through interference with actin-1/profilin dynamics"

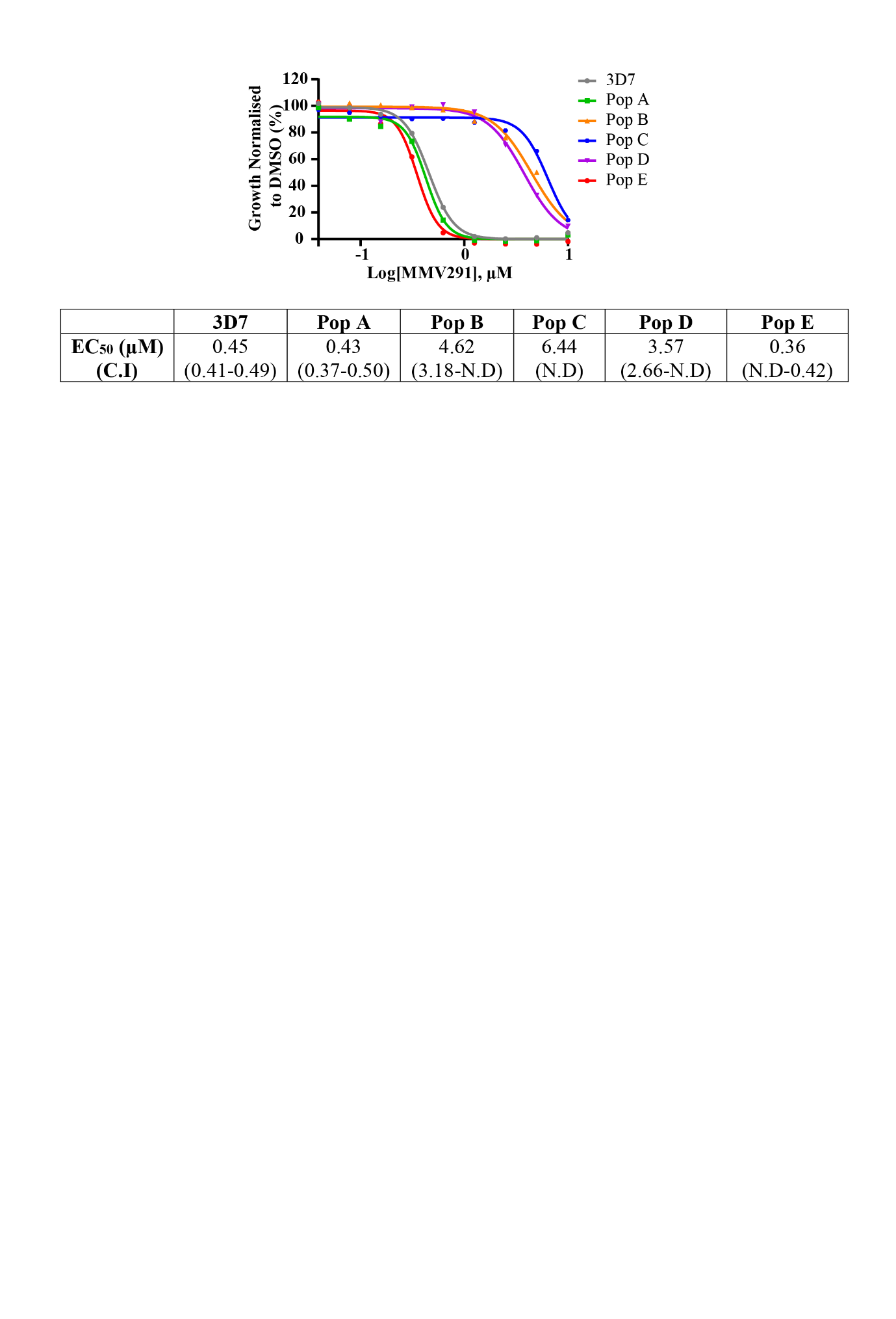

**Figure S1. *In vitro* resistance to MMV291**

Viable parasites recovered after three rounds of drug cycling were tested against a titration of MMV291 in a 72-hour lactate dehydrogenase (LDH) growth assay. Parasite growth was normalised to parasite’s grown in 0.1% DMSO which indicated three resistant populations were obtained (B, C and D) with an 8 to 14-fold increase in EC_50_ compared to 3D7. Data points represent the average of three technical replicates. C.I indicates 95% confidence intervals for EC_50_ values which were derived from nonlinear regression curves in GraphPad Prism. N.D= not determined.

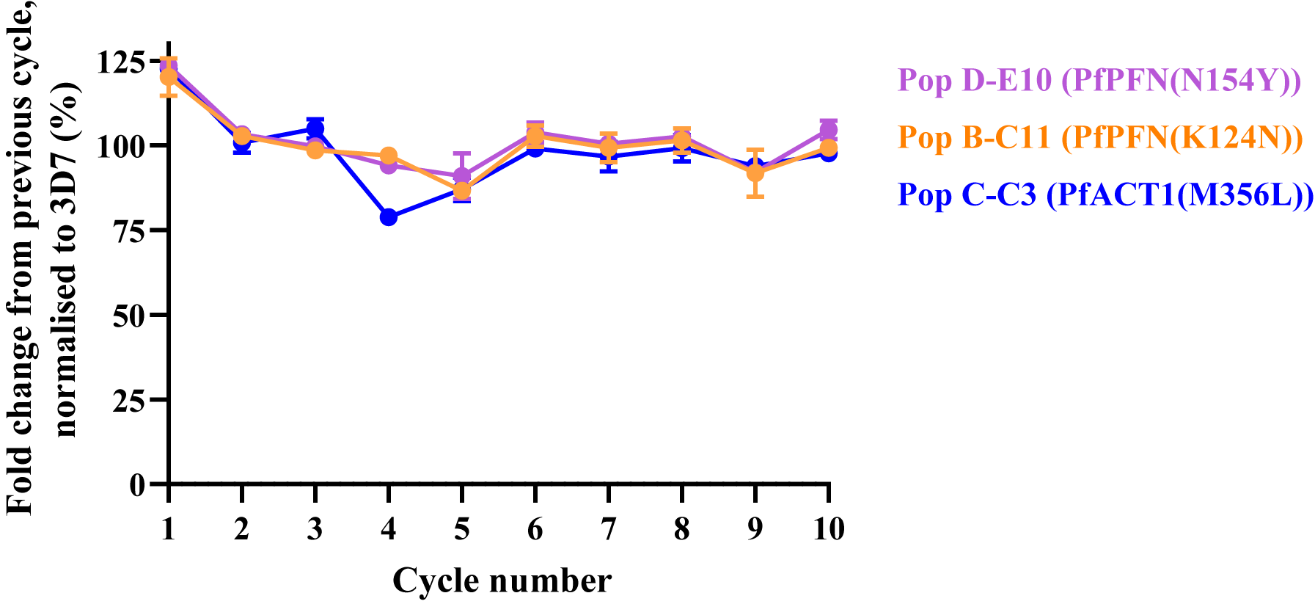

**Figure S2. MMV291 resistant lines do not have reduced parasite fitness**

The growth of three MMV291 resistant population clones, Pop D-E10, Pop B-C11 and Pop C-C3, with the corresponding PFN(N154Y), PFN(K124N) and ACT1(M356L) mutations, along with 3D7 wildtype parasites, were compared in a 10-cycle growth assay. During each cycle, an aliquot of culture was harvested from each parasite line and frozen until completion of the assay whereby parasite lactate dehydrogenase was measured as a marker for parasite growth. The fold change in parasitemia was calculated from the previous cycle for each parasite line, which was then expressed as a percentage of the 3D7 fold change. This demonstrated that there was no comparative growth defect associated with the resistant lines, indicating that the mutations in PfPFN and PfACT1 did not reduce the fitness of these parasites. Error bars represents the standard deviation from one experiment comprising of three technical replicates.

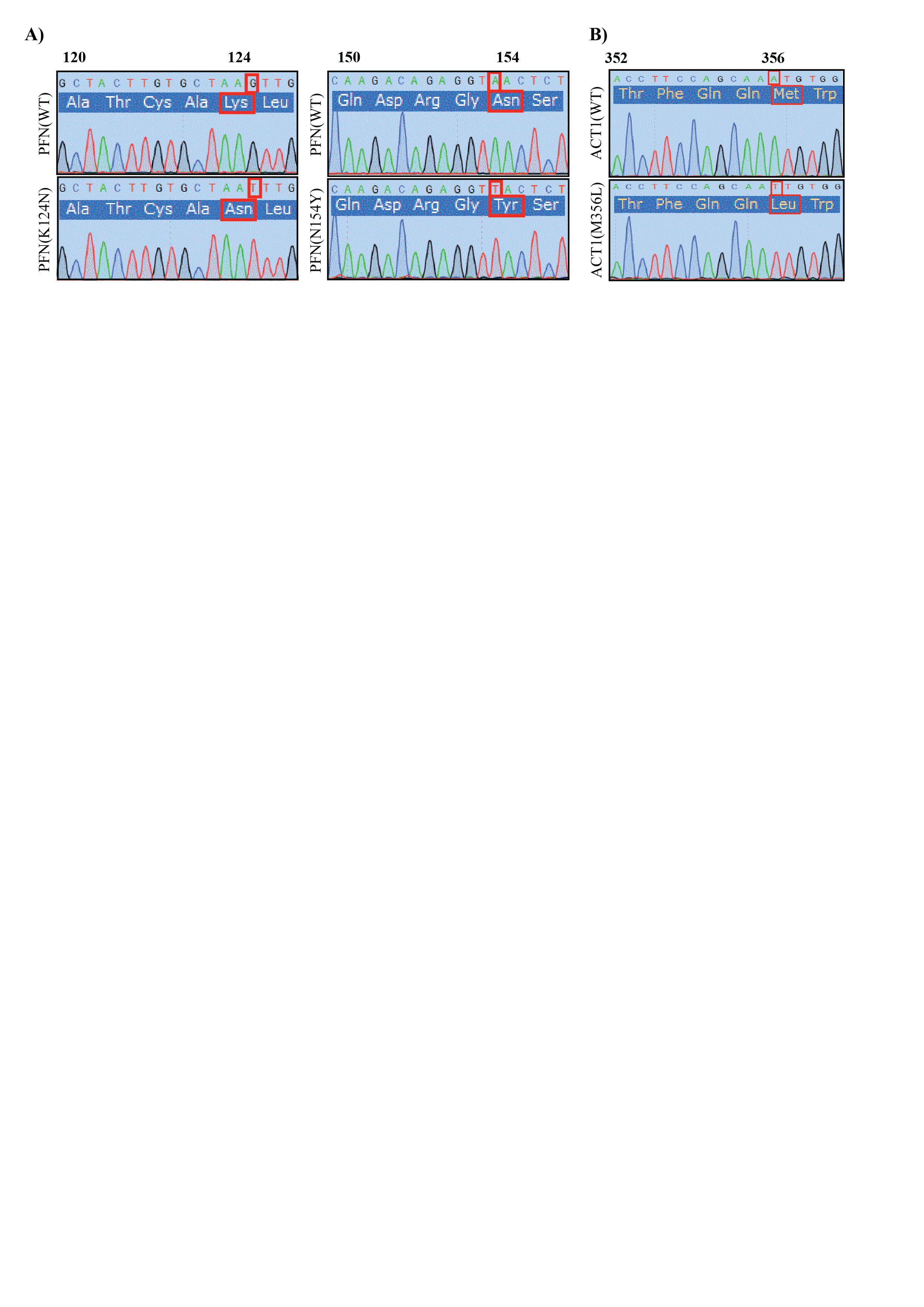

**Figure S3. Chromatograms from integrated parasites containing the MMV291-resistant alleles**

The products produced from diagnostic PCRs were sequenced and the resistant mutations were confirmed to be present for **(A)** K124N (AAG-AAT) and N154Y (AAC-TAC) in *profilin* and **(B)** M356L (ATG-TTG) in *actin-1*.

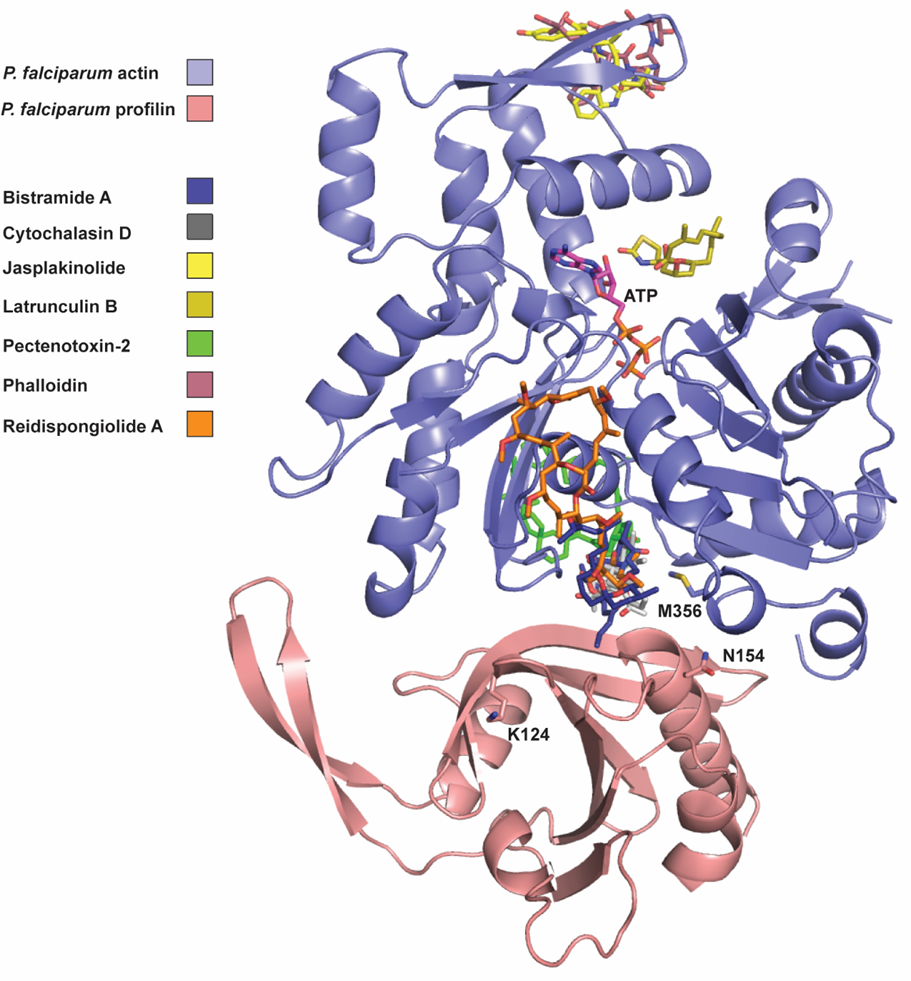

**Figure S4. Model of *P. falciparum* profilin and actin-1 with known actin binders**

*P. falciparum* profilin (pink) (PDB: 2JKG) ((Kursula et al., 2008), structure 16: 1638) and actin-1 (blue) (ATP, magenta) (PDB: 6I4E) (Kumpula et al., 2019) heterodimeric complex showing regions of the proteins where actin inhibitors are known to bind relative to the MMV291 *P. falciparum* mutations. The actin inhibitors aligned to *P. falciparum* actin-1 and shown are Bistramide A (blue) (aligned from *O. cuniculus* actin, PDB: 2FXU) (Rizvi et al., 2006), Cytochalasin D (grey) (aligned from *D. melanogaster* actin, PDB: 3EKU) (Nair et al., 2008), Jasplakinolide (yellow) (aligned from *P. falciparum* F-actin, PDB: 5OGW) (Pospich et al., 2017), Latrunculin B (gold) and Pectenotoxin-2 (green) (aligned from *O. cuniculus* actin, PDB: 2Q0U) (Allingham et al., 2007), Phalloidin (maroon) (aligned from *G. gallus* F-actin, PDB: 7BTI) (Kumari et al., 2020), Reidispongiolide A (orange) (aligned from *O. cuniculus* actin PDB: 2ASM) (Allingham et al., 2005).

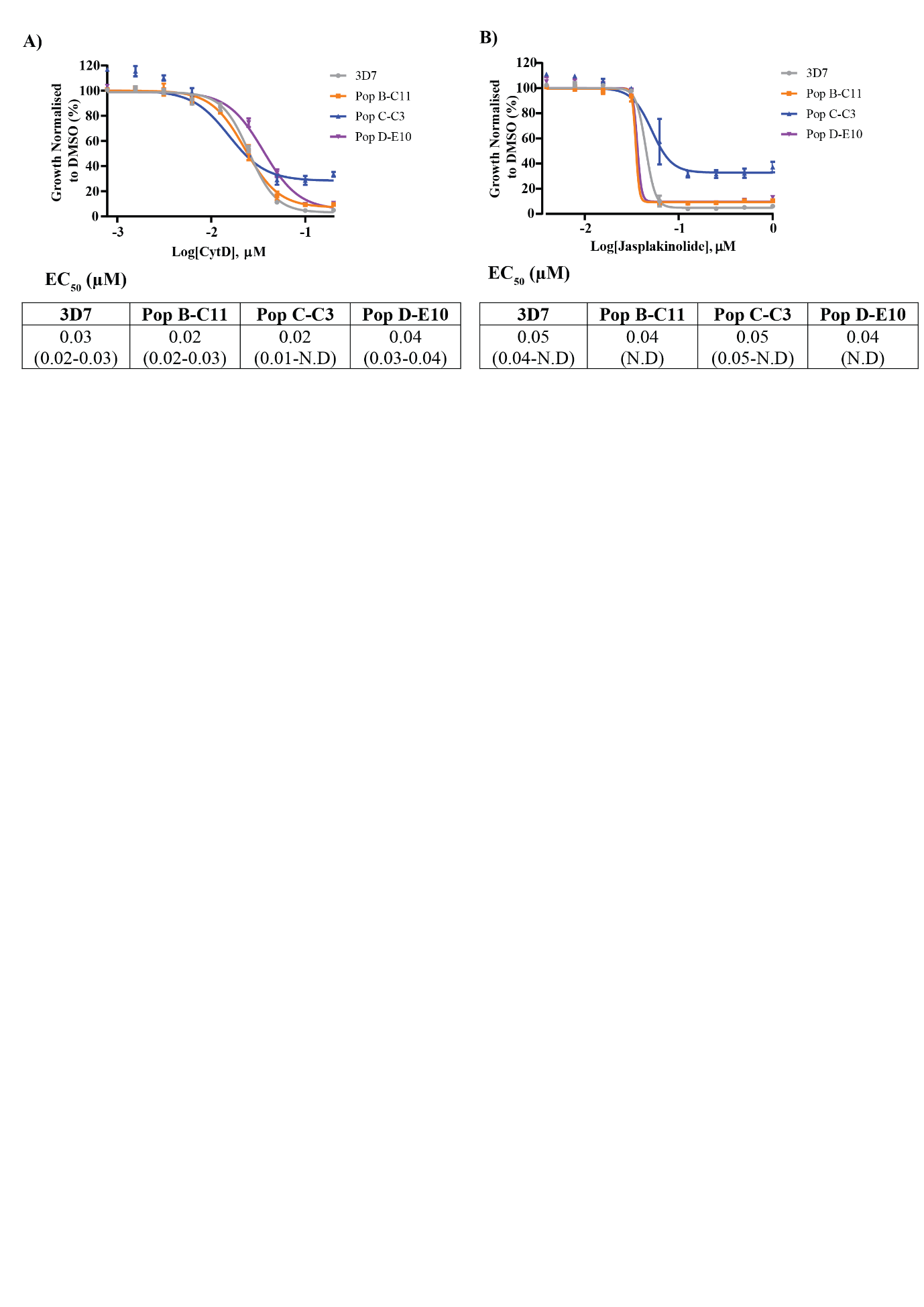

**Figure S5. MMV291 resistant lines are not cross-resistant to other actin polymerisation inhibitors**

A titration of the actin polymerisation inhibitor, Cytochalasin D (CytD) **(A)**, and actin polymerisation stabiliser, Jasplakinolide **(B)**, were tested against the MMV291-resistant lines and 3D7 parasites in a 72-hour growth assay. This revealed that MMV291-resistant parasites did not exhibit cross resistance to CytD and Jasplakinolide, indicating that MMV291 has an alternate mechanism of action. Growth has been normalised to that of parasites grown in 0.1% DMSO and error bars represent the standard deviation of three biological replicates. EC_50_ values were derived from nonlinear regression curves in GraphPad Prism with 95% confidence intervals of these values specified in brackets.

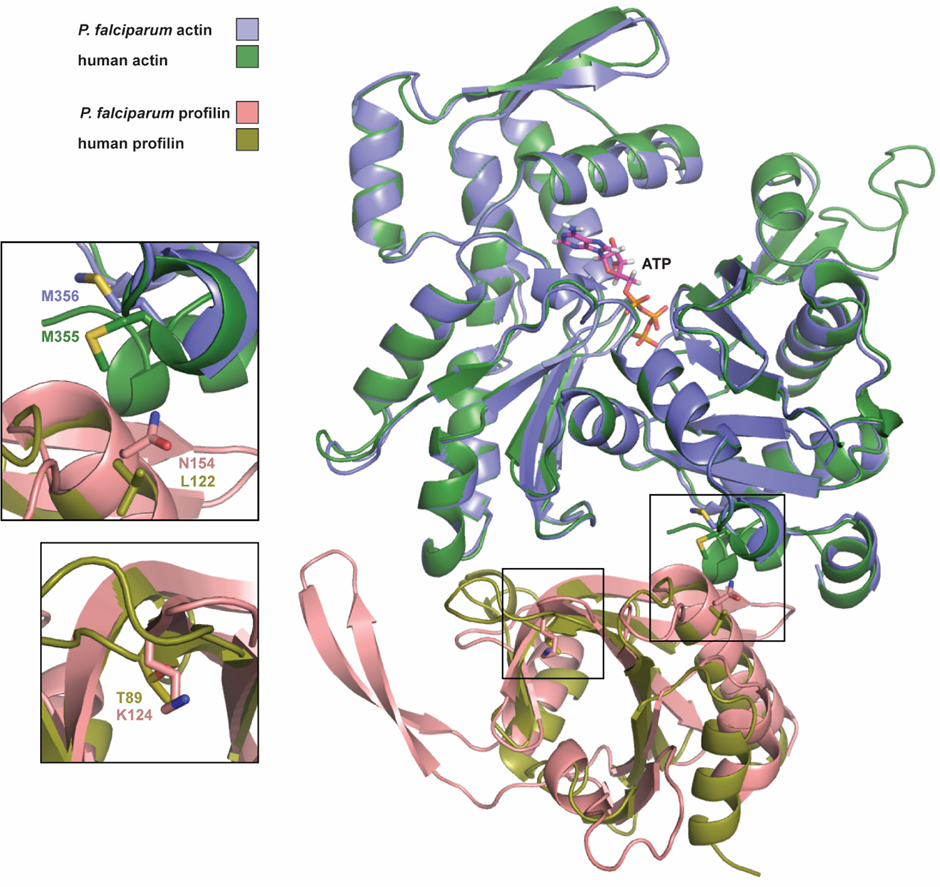
**Figure S6. A model of the comparison between mutation locations in human and *P. falciparum***

An X-ray structure human profilin (gold) (PDB: 2PBD) and a homology model of human actin (green) (created by SWISS-MODEL (Waterhouse et al., 2018) using *O. cuniculus* actin (PDB: 2PBD) (Ferron et al., 2007) aligned with *P. falciparum* profilin (pink) (PDB: 2JKG) (Kursula et al., 2008) and actin (blue) (PDB: 6I4E) (Kumpula et al., 2019) showing the similarity of the heterodimeric complex. The positions of the MMV291 *P. falciparum* mutations and the associated human amino acids are shown for comparison.

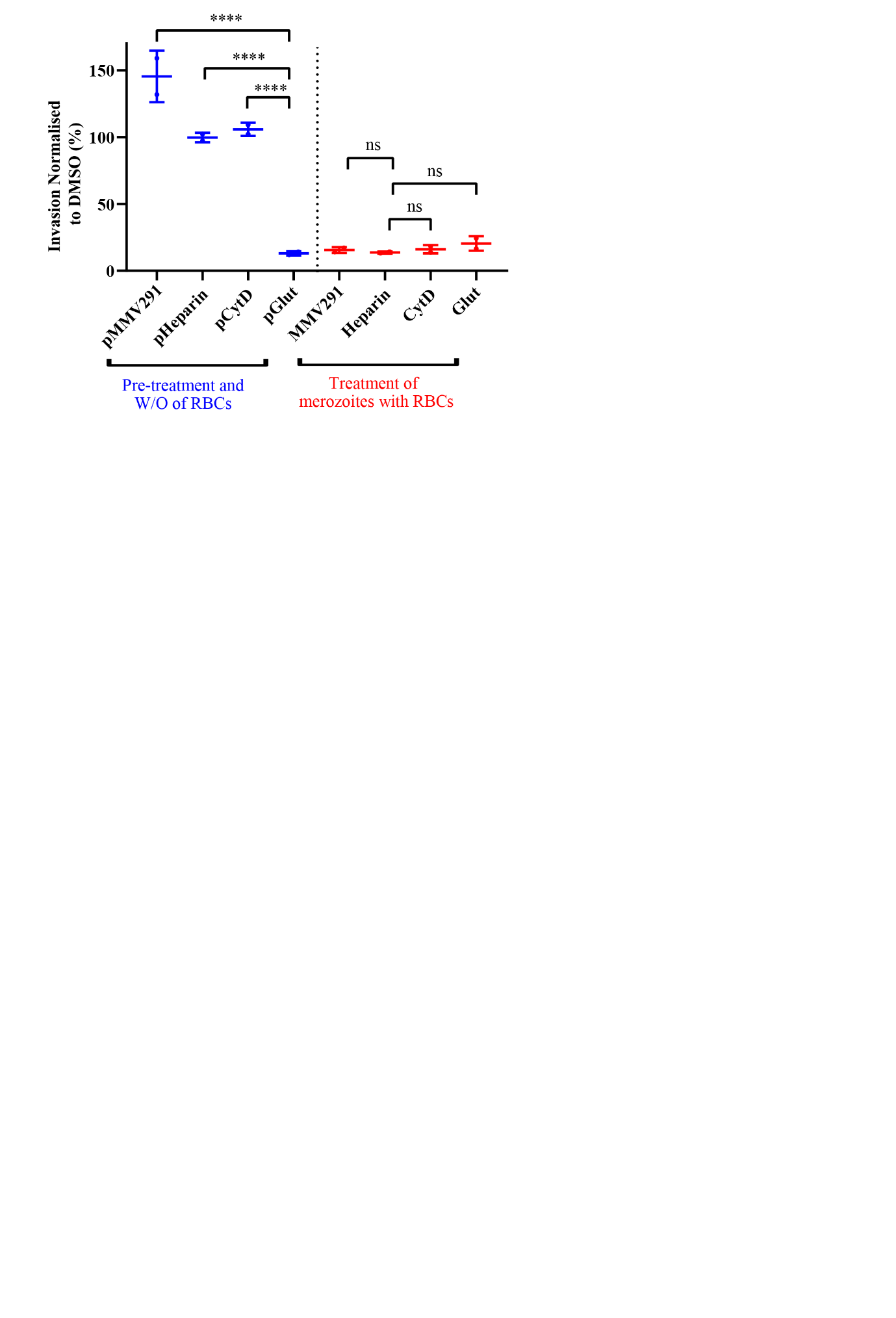

**Figure S7. MMV291 pre-treatment of uninfected RBCs does not inhibit merozoite invasion**

Uninfected RBCs were pre-treated with invasion inhibitory compounds (blue); 10 µM MMV291, 100 µg/mL heparin, 2 μM cytochalasin D (CytD), or 0.0025% glutaraldehyde (Glut) for 30 min at 37°C, after which the cells were washed (W/O) to remove the inhibitors. Purified merozoites were then allowed to invade the pre-treated RBCs. In parallel, merozoites were added to untreated RBCs in the presence of these inhibitors (red). After incubation for 30 minutes at 37°C, the compounds were washed out and parasites allowed to grow for 24 hours. Successful invasion was assessed by measuring the bioluminescence levels of trophozoite stage parasites expressing a nanoluciferase reporter and invasion rate was normalised to the DMSO vehicle control. This demonstrated that unlike the fixative glutaraldehyde, pre-treatment with MMV291 did not reduce merozoite invasion of RBCs, producing a similar profile to the invasion inhibitory molecules, heparin and CytD. Error bars represent the standard deviation of two biological replicates with statistical analyses performed in GraphPad Prism using a one-way ANOVA with pre-treated RBCs compared to glutaraldehyde (blue) and merozoite treatment compared to heparin (red). **** indicates P<0.0001, ns indicates not significant (P>0.05).

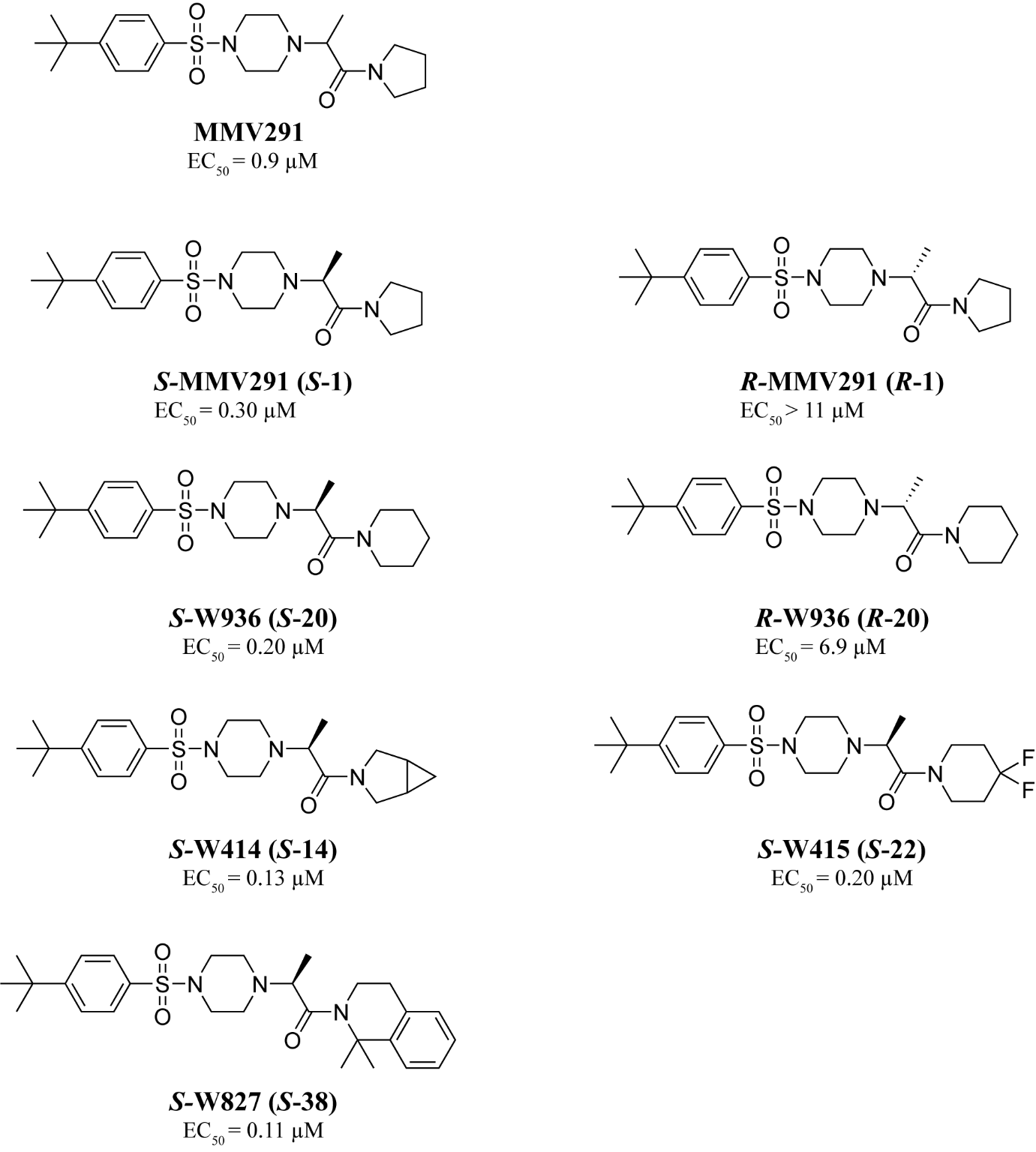

**Figure S8. MMV291 analogue structures**

The chemical structures and corresponding EC_50_ values against the RBC stage of *P. falciparum* used in this study with original compound names from Nguyen, *et al*. (2021) specified in brackets.

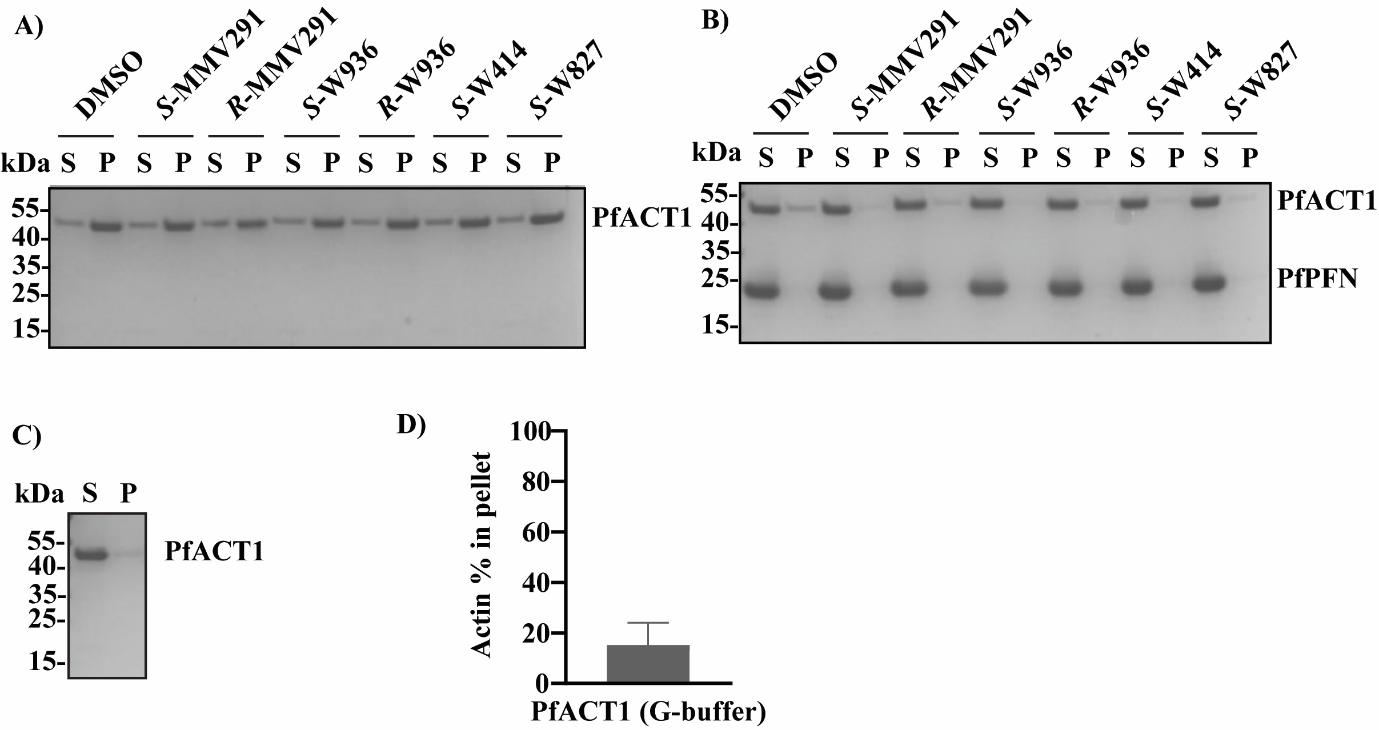

**Figure S9. Actin sedimentation assay gels and quantification of the actin G-buffer control**

Sedimentation samples consisting of 4 µM PfACT1 **(A)** and with 16 µM PfPFN **(B)** in the presence of 25 µM MMV291 analogues or DMSO as well as 4 µM PfACT1 alone in G-buffer **(C)** were analysed on 4-20% Mini-PROTEAN TGX gels and visualized with PageBlue stain. S denotes supernatant and P pellet. **D)** Quantification of the relative amount of actin in the pellet fraction for PfACT1 in G-buffer whereby 15 ± 9 % of PfACT1 sedimented to the pellet fraction. Results are reported as mean ± standard deviation. The data are based on at least three independent assays each performed in triplicate, with a representative gel presented.

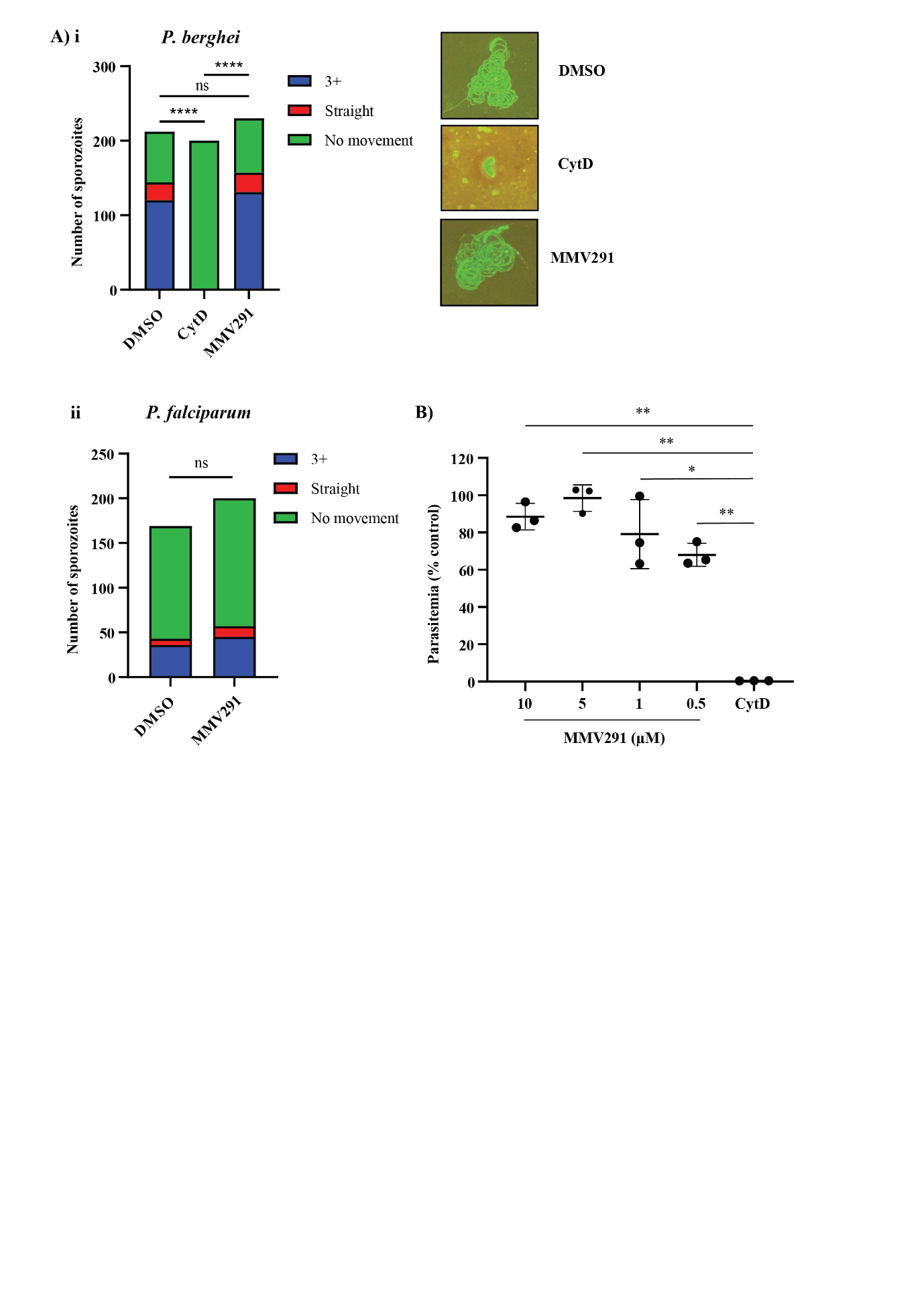

**Figure S10. MMV291 has no effect on sporozoite motility or invasion**

**A)** Sporozoites expressing GFP were used to measure motility via the quantification of fluorescent trails. This revealed that similarly to DMSO, MMV291 had no effect on sporozoite motility in *P. berghei* **(i)** or *P. falciparum* **(ii)**, whilst cytochalasin D (CytD) significantly reduced motility. 3+ indicates 3 or more trails observed. **B)** *In vitro* human liver cells were incubated with a titration of MMV291 in the presence of 20 000 sporozoites expressing a luciferase protein. After 52 hours, cells were lysed and luciferase activity was measured to correlate with sporozoite invasion rate. In contrast with CytD (10 µM) treatment, MMV291 did not reduce invasion rate of sporozoites at concentrations tested. Error bars represent the standard deviation across three biological replicates each comprised of three technical replicates. Statistical analysis performed via a Chi-square (A) and unpaired t-test (B) using GraphPad Prism. * P<0.05, ** P<0.01, **** P<0.0001, ns indicates not significant.

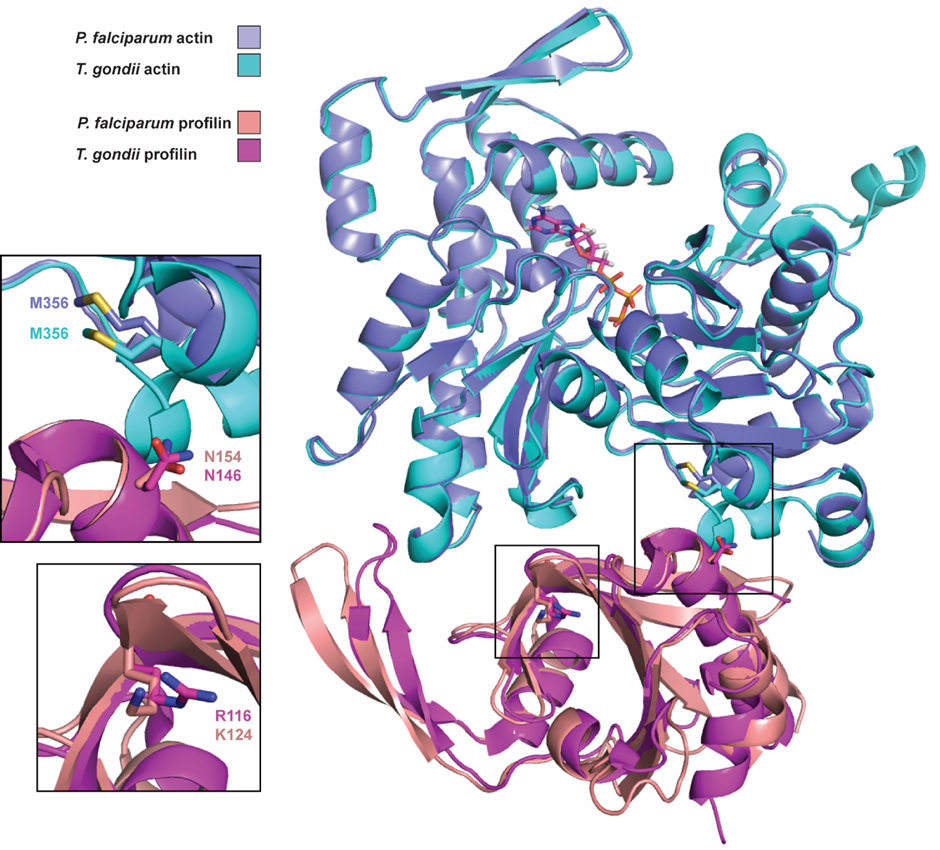

**Figure S11. A model of the comparison between mutation locations in *T. gondii* and *P. falciparum***

*T. gondii* profilin (magenta) and actin (cyan) aligned with *P. falciparum* profilin (pink) and actin (blue) showing the similarity of the heterodimeric complex and the positions of the MMV291 *P. falciparum* mutations. The X-ray structure of *T. gondii* profilin (PDB: 3NEC) (Kucera et al., 2010) and a homology model of *T. gondii* actin (created by SWISS-MODEL (Waterhouse et al., 2018) using the X-ray structure of *P. falciparum* actin (PDB: 6I4K) (Kumpula et al., 2019) were used to create the model. The X-ray structure of *O. cuniculus* actin and human profilin (PDB: 2PBD) (Ferron et al., 2007) was utilised as a template to spatially overlay the *P. falciparum* actin and profilin in the heterodimer model.

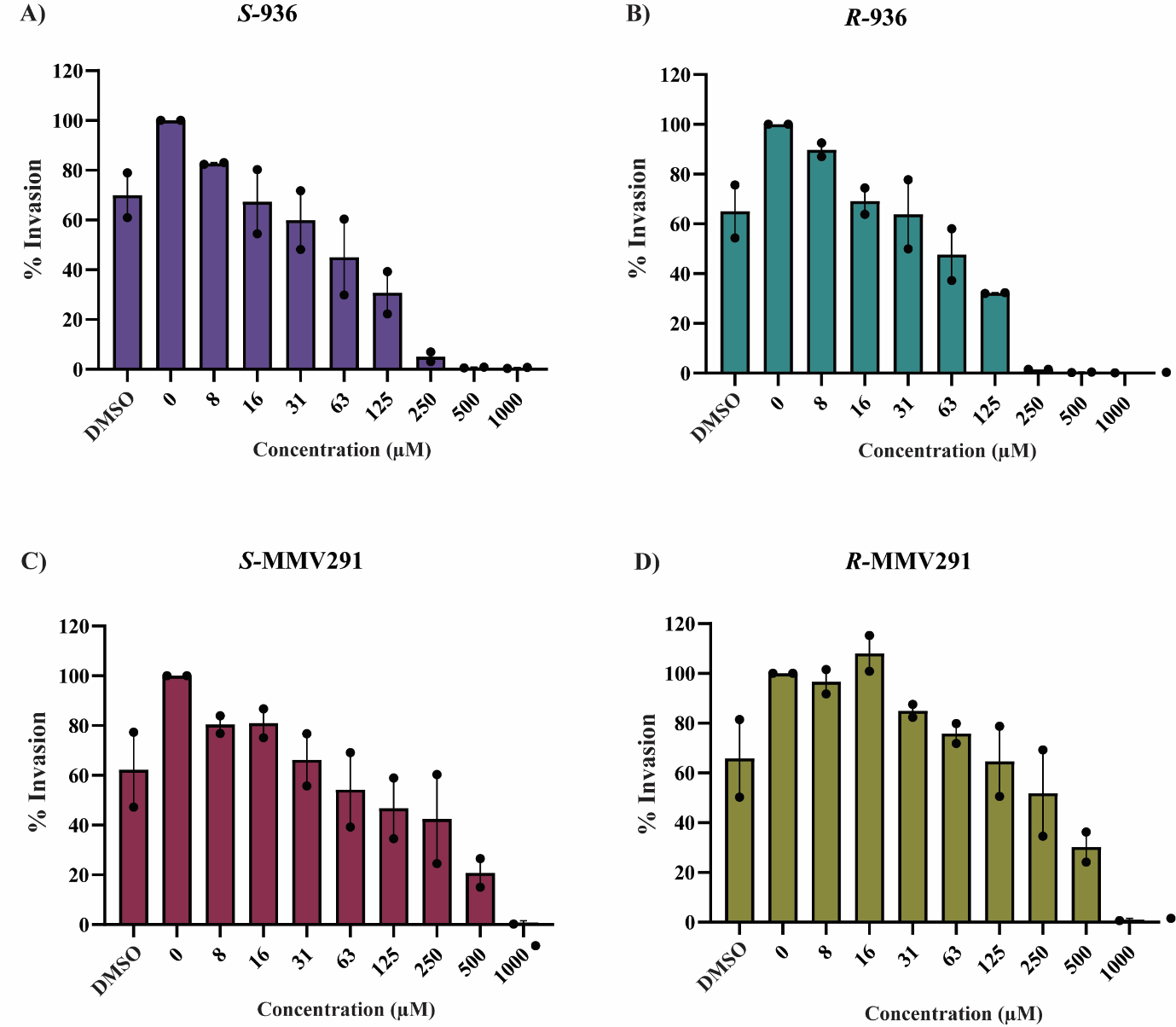

**Figure S12. MMV291 series show limited activity against *T. gondii* invasion**

Nanoluciferase expressing parasites were liberated from their host cell and incubated with the MMV291 analogues before being added back to fibroblasts and allowed to invade for 1 hour before compounds were washed out. After a 24-hour incubation, cells were then lysed and the relative light units was quantified to correlated with *T. gondii* invasion rate. This showed MMV291 analogues *S*-W936 **(A)**, *R*-W936 **(B)**, *S*-MMV291 **(C)** and *R*-MMV291 **(D)** had some inhibitory activity against invasion at high concentrations. DMSO was included to the same amount as the highest concentration of analogue to account for DMSO-related effects (30% reduction in invasion). Error bars indicate the standard deviation from two biological repeats.

Table S1. Primers used to construct donor plasmids

| **To knock-in mutations in Profilin** | | | |
| --- | --- | --- | --- |
| HR1A | PFN_BglII.1F (a) | *AGATCT*TTCATGGGACAGTTATTTAAATGATCGCCT | *Bgl*II |
|  | PFN_2R (b) | AGATAGTTTGACCTTCGTTGATCGTTTTGGTAGTTTTAGTACCA |  |
| HR1B | PFN_CO.3F (c) | CAACGAAGGTCAAACTATCTTGGTTGT |  |
|  | PFN_CO.Spe.4R (d) | *ACTAGT*CTATTGAGAAGATTCAGCCAATTCCTTAGCGA | *Spe*I |
| HR2 | PFN_EcoRI.5F (e) | *GAATTC*TGGTCGTTTTTAATGAAGGATATGCTCCTGA | *EcoR*I |
|  | PFN_KasI.6R (f) | *GGCGCC*CTATTGACTGCTTTCAGCTAGCTCTTTTGC | *Kas*I |
| gRNA | PFN_gRNA | AAATGAAGGACAAACGATCC/TGG |  |
| Introduce K124N | PFN_K12N_F (g) | TACTTGTGCTAATTTGAAGGGTGGT |  |
|  | PFN_K124N_R (h) | ACCACCCTTCAAATTAGCACAAGTA |  |
| Confirm integration | PFN_5'UTR_F (i) | CCTTTTCTTTTTCTTTCTTCCTCTCACTCTCTCATACTCTCT |  |
|  | PFN_WT_R (j) | TGGAGTTGTCTATGCTTGTGTAGCTCAGGG |  |
|  | glmS_R (k) | AGATCATGTGATTTCTCTTTGTTCA |  |
| **To knock-in mutations in Actin-1** | | | |
| HR1A | ACT_BglII.1F (l) | *AGATCT*AAAATGGGAGAAGAAGATGTTCAAGCT | *Bgl*II |
|  | ACT_2R (m) | GTCGAGACGCATAATCGCGTGTGGTAAAGCATAACCTTCATAAATTGGA |  |
| HR1B | ACT_CO.3F (n) | CACGCGATTATGCGTCTCGAC |  |
|  | ACT_CO.Spe.4R (o) | *ACTAGT*TTAGAAGCATTTACGATGGACAATAGA | *Spe*I |
| HR2 | ACT_EcoRI.5F (p) | *GAATTC*TTTAGCTGGTAGAGATTTAACTGAATATTTAATGA | *EcoR*I |
|  | ACT_KasI.6R (q) | *GGCGCC*TTAGAAACATTTTCTGTGGACAATACTTGGTCCT | *Kas*I |
| gRNA | ACT_sgRNA | TCTAATCTCATAATTGCATG/TGG |  |
| Introduce M356L | ACT_CO_M356L.F (r) | CTCTTTCCACCTTCCAGCAATTGTGGATTACT |  |
|  | ACT_CO_M356L.R (s) | AGTAATCCACAATTGCTGGAAGGTGGAAAGAG |  |
| Replace internal *Bgl*II site | ACT_3'_repairF (t) | GGATATTCGTAAAGACCTTTATGGAAATATCGT |  |
|  | ACT_5'_repairR (u) | ACGATATTTCCATAAAGGTCTTTACGAATATCC |  |
| Confirm integration | ACT_5'UTR_F (v) | CCATTTGGTGATTAGTTTTTACTGAC |  |
|  | ACT_WT_R (w) | GCTCCAGAAGAACACCCAGTGTTATTAAC |  |
